# Distinct cellular mediators drive the Janus Faces of Toll-like Receptor 4 regulation of network excitability which impacts working memory performance after brain Injury

**DOI:** 10.1101/750869

**Authors:** Akshata A. Korgaonkar, Ying Li, Susan Nguyen, Jenieve Guevarra, Kevin C H Pang, Vijayalakshmi Santhakumar

## Abstract

The mechanisms by which the neurophysiological and inflammatory responses to brain injury contribute to memory impairments are not fully understood. Recently, we reported that the innate immune receptor, toll-like receptor 4 (TLR4) enhances AMPA receptor (AMPAR) currents and excitability in the dentate gyrus after fluid percussion brain injury (FPI) while limiting excitability in controls. Here we examine the cellular mediators underlying TLR4 regulation of dentate excitability and its impact on memory performance. In *ex vivo* slices, astrocytic and microglial metabolic inhibitors selectively abolished TLR4 antagonist modulation of excitability in controls, without impacting FPI rats, demonstrating that glial signaling contributes to TLR4 regulation of excitability in controls. In glia-depleted neuronal cultures from naïve mice, TLR4 ligands bidirectionally modulated AMPAR charge transfer demonstrating the ability of neuronal TLR4 to regulate excitability, as observed after brain injury. *In vivo* TLR4 antagonism reduced early post-injury increases in mediators of MyD88-dependent and independent TLR4 signaling without altering expression in controls. Blocking TNFα, a downstream effector of TLT4, mimicked effects of TLR4 antagonist and occluded TLR4 agonist modulation of excitability in slices from both control and FPI rats. Functionally, transiently blocking TLR4 *in vivo* improved impairments in working memory observed one week and one month after FPI, while the same treatment impaired memory function in uninjured controls. Together these data identify that distinct cellular signaling mechanisms converge on TNFα to mediate TLR4 modulation of network excitability in the uninjured and injured brain and demonstrate a role for TLR4 in regulation of working memory function.

**Highlights:**

- TLR4 suppresses dentate excitability in controls through signaling involving glia
- Neuronal TLR4 signaling underlies enhanced dentate excitability after brain injury
- TNFα contributes to TLR4 regulation of excitability in the injured brain
- Altering TLR4 signaling impacts working memory performance
- TLR4 signaling is a potential target to improve working memory after brain trauma

## 1.1 Introduction

A growing number of studies suggest that immune signaling can modulate the nervous system’s function and plasticity (Pribiag and Stellwagen, 2014; van Vliet et al., 2018). Activation of glial inflammatory signals including TNFα and purinergic receptors have been shown to impact synaptic plasticity and hippocampal memory function (Beattie et al., 2002; Belarbi et al., 2012; Pascual et al., 2012; Pribiag and Stellwagen, 2014). Increases in neuronal excitability and immune activation are hallmarks of traumatic brain injury and present a condition in which neuroimmune interactions are potentially accentuated (Chiu et al., 2016; Neuberger et al., 2017a). While post-traumatic inflammatory responses are implicated in neurodegeneration, and immunosuppressants can improve neurological outcomes after brain injury (Saletti et al., 2019), mechanisms underlying immune regulation of neuronal function and behaviors are not fully understood. Among the immune pathways activated by brain injury, Toll-like receptor 4 (TLR4), a member of the innate immune Pattern Recognition Receptors, has been shown to contribute to sterile inflammatory responses (Laird et al., 2014; Vezzani et al., 2011; Zhu et al., 2014). Although TLR4 is traditionally recruited in defense against by microbial pathogens, it can also be activated by endogenous ligands released from injured tissue and dying cells making it a critical link between brain injury and the ensuing inflammatory response (Kielian, 2006). While TLR4 activation after brain injury is well documented (Ahmad et al., 2013; Laird et al., 2014; Ye et al., 2014), we recently demonstrated the preferential localization of TLR4 in hippocampal dentate granule cell layer and hilar neurons rather than glia (Li et al., 2015). Moreover, we identified that TLR4 signaling reduced excitability in *ex vivo* slices from sham injured animals while enhancing excitability after brain injury (Li et al., 2015). The cellular mechanisms and functional consequences of this differential TLR4 regulation of dentate network excitability are currently unknown.

TLR4 is known to be expressed in both neurons and glia (Okun et al., 2011). However, a majority of the established functional effects of TLR4 in the CNS involve glial signaling and little is known about the role of neuronal TLR4. Consequently, signaling downstream of TLR4 has been elucidated primarily in glia. Activation of microglial and astrocytic TLR4 is known to engage two distinct signaling cascades, the pathway dependent on the Myeloid Differentiation factor 88 (MyD88) and the MyD88-independent pathway which activates the TIR-domain-containing adapter-inducing interferon-β (TRIF) (Kawai and Akira, 2007). Recruitment of the MyD88-dependent pathway in astrocytes and microglia leads to transcription of nuclear factor-kappa B (NF-κB) and production of the pro-inflammatory cytokine Tumor Necrosis Factor α (TNFα). Glia-derived TNFα has been shown to activate neuronal TNF receptor 1 (TNFR1) and mediate increases in NMDA receptor-dependent synaptic plasticity and AMPA receptor (AMPAR) surface expression (Stellwagen et al., 2005). The MyD88-independent pathway involves the recruitment of TRIF to activate Interferon regulatory factor 3 (IRF3), which in turn causes the production of interferon-β. Whether these or other signaling pathways are recruited by neuronal TLR4 remains unknown.

A second aspect of interest concerns the functional consequences of TLR4 regulation in hippocampal dentate excitability. Sparse granule cell activity is critical for dentate spatial working memory function (Dengler and Coulter, 2016). Conditions that enhance dentate excitability or impair inhibition compromise memory function. Indeed, brain injury leads to both increased dentate excitability and impairments in memory performance (Gupta et al., 2012; Hamm et al., 1996; Santhakumar et al., 2001; Semple et al., 2018). Several lines of evidence suggest a role for TLR4 in hippocampal memory processing. Mice lacking TLR4 show enhanced memory processing and mutations that alter TLR4 signaling impact hippocampal long-term potentiation and spatial reference memory (Costello et al., 2011; Okun et al., 2012). Changes in hippocampal memory processing in mice lacking TLR4 have been associated with increases in cells expressing the immediate early gene Arc, which is consistent with enhanced excitability. These findings suggest that TLR4 modulation of dentate excitability could impact memory performance (Jeltsch et al., 2001; Xavier et al., 1999). The current study was conducted to determine the signaling mechanisms underlying TLR4 regulation of network excitability in the injured brain and its impact on behavioral outcomes.

## 2. Materials and Methods

All procedures were performed under protocols approved by the Institutional Animal Care and Use Committee of the Rutgers New Jersey Medical School, Newark, New Jersey and are consistent with the ARRIVE guidelines.

### 2.1 Fluid percussion injury

Juvenile male Wistar rats (25-27 days old) were subject to moderate (2-2.2 atm) lateral fluid percussion injury (FPI) or sham-injury using standard methods (Li et al., 2015; Neuberger et al., 2017b). Studies were restricted to males to avoid confounds due to cyclical changes in immune responses and behavioral effects of injury in females (Potter et al., 2019; Roof and Hall, 2000). Briefly, under ketamine (80mg/kg)-xylazine (10mg/kg) anesthesia (i.p.), rats underwent stereotaxic craniotomy (3mm dia, -3mm bregma, and 3.5mm lateral to sagittal suture) to expose the dura and bond a Luer-Lock syringe hub. The next day, randomly selected animals received a brief (20ms) impact on the intact dura using the FPI device (Virginia Commonwealth University, VA) under isoflurane anesthesia. Sham rats underwent all procedures except delivery of the pressure wave. Animals with implants dislodged during impact and those with <10 sec apnea following injury or injuries in which the pressure waveforms were jagged were excluded.

### 2.2 Field Electrophysiology

One week after FPI or sham-injury rats were euthanized under isoflurane and horizontal brain slices (400µm) were prepared in ice-cold sucrose-artificial cerebrospinal fluid (ACSF) containing (in mM) 85 NaCl, 75 sucrose, 24 NaHCO_3_, 25 glucose, 4 MgCl_2_, 2.5 KCl, 1.25 NaH_2_PO_4_, and 0.5 CaCl_2_ (Li et al., 2015; Yu et al., 2016). Recordings were restricted to the side of injury (ipsilateral).

Field recordings were obtained in an interface recording chamber (BSC2, Automate Scientific, Berkeley, CA) perfused with ACSF containing (in mM) 126 NaCl, 2.5 KCl, 2 CaCl2, 2 MgCl2, 1.25 NaH2PO4, 26 NaHCO3 and 10 D-glucose at 32–33°C. Granule cell population responses were recorded using patch pipettes with ACSF, in response to stimuli delivered through bipolar tungsten stimulating electrodes in the perforant path. Population spike amplitude was measured as the amplitude of the first negative deflection overriding the field EPSP waveform, as described previously (Neuberger et al., 2014). In all drug incubation experiments, pre and post drug recordings were conducted in the same slices with a 45 min incubation between recordings. Slices were recorded in control ACSF and transferred to incubation chambers containing ACSF or one of the following drug cocktails (a) CLI-095, (b) CLI-095 + fluoroacetate + minocycline (c) fluoroacetate + minocycline (d) HMGB1 (e) anti-TNFα or (f) HMGB1 + anti-TNFα for 45 minutes before returning slices back to the recording chamber. The drug were used at the following concentrations: CLI-095, 10ng/ml; fluoroacetate, 1mM; minocycline, 50nM; HMGB1, 10 ng/ml and anti-TNFα, 1µg/ml. Electrode positions were maintained during pre and post incubation recordings. Control incubations in ACSF between recordings confirmed that perforant path-evoked dentate population spike amplitude was stable during experimental manipulations (Suppl. Fig. 1). All electrophysiology data were low pass filtered at 3 kHz, digitized using DigiData 1440A, acquired using pClamp10 at 10 kHz sampling frequency, and analyzed using ClampFit 10. In experiments in which the percent change in population spike was analyzed, data from recordings in which no population spikes were observed in ACSF were excluded from analysis.

### 2.3 Neuronal Cultures

Primary hippocampal neurons were obtained from the hippocampi of E17–18 C57Bl/6J mouse embryos of both sexes. Hippocampi were transferred to 15ml tissue culture tubes and the volume was adjusted to 10ml with Hanks balanced salt solution (HBSS) and were then gently centrifuged at 1000 rpm for 1 min. HBSS was then removed from the tube and tissue was digested in 5ml of 0.25% trypsin and 75µl of DNase (200Units/mg) solution for 20 min at 37°C. Trypsin/DNase mix was then removed from tissue and the tissue was washed with characterized fetal bovine serum. Tissue was transferred to Neurobasal plating media-A (NB-A) and mechanically dissociated by trituration with a Pasteur pipet. Cells were pelleted by centrifugation at 1500 rpm for 5 min. Cells were taken up in 20µl of NB-A plating medium and counted in a hemocytometer. Approximately 0.5-1.0 x 10^6^ cells per well were plated on Poly-D-Lysine (PDL) coated 15mm coverslips in NB-A medium containing B27 and L-glutamine and cultured at 37°C in 5% C02/95% air. Medium was changed every 3-4 days. Ara-C was added to media on day 3 for complete removal of glia. Cultured coverslips were stained for GFAP and IBA-1 for confirmation of complete absence of glia. 11- to 13-day old cultures were used for physiological recordings. Recordings were obtained from independent culture plates obtained from 3-4 pregnant dams.

Cultured neurons were visualized under IR-DIC using a Nikon Eclipse FN-1 microscope and a 40X, 0.8 NA water-immersion objective. Whole cell recordings were obtained using MultiClamp 700B (Molecular Devices) at 32–33°C. Voltage clamp recordings of AMPAR currents were performed using borosilicate microelectrodes (4–6 MΩ) containing (in mM) 140 Cs-gluconate, 10 HEPES, 2 MgCl2, 0.2 EGTA, 2 Na-ATP, 0.5 Na-GTP, 10 phosphocreatine and 0.2% biocytin in the presence of SR95531 (10µM) to block GABAA receptors and D-APV (50µM) to block NMDA receptors. Recordings were rejected if the cell had more than a 20% change in series resistance over time.

### 2.4 Drug administration

One day after injury, a randomly assigned cohort of FPI and sham rats received either vehicle (saline) or a synthetic TLR4 antagonist, CLI-095 (0.5mg/kg, s.c.) for 3 days starting 24 hrs. after injury. A second group underwent stereotaxic injection (Hamilton syringe-26 G) of 5µl of saline or the highly selective TLR4 antagonist LPS-RS Ultrapure (LPS-RS*U*, 2mg/ml) in the hippocampus on the side of injury. Injections were delivered through the implanted syringe hub (AP: 3.0mm, ML: 3.5mm, DV: 3.2mm). Drugs were delivered one day after injury at a rate of 1µl/5 mins under isoflurane anesthesia (2% in 95% O2, 5% CO2) delivered through a nose cone.

### 2.5 Western blotting

Western blots of protein from fresh hippocampal tissue were obtained from rats perfused with cold ACSF (4°C) 3 days after FPI or sham-injury as described previously (Li et al., 2015) using antibodies listed in Table 1. Protein concentration of the sample lysates was measured using BCA assay (Santa Cruz). Equal amounts of protein samples were diluted at a ratio of 1:1 in Laemmli sample buffer (Sigma) and separated on pre-cast gel (4–12% Tris–glycineBio-Rad). Chemiluminescent detection was performed with ECL western blotting detection reagent (Westdura, Thermo Scientific) using FluoroChem 8800. Densitometric quantification was determined using Image-J software (NIH) and normalized to β-actin density.

**Table 1.** Summary Table of Antibodies used in the study.

### 2.6 Spatial Working Memory test

A delayed match to position working memory task was conducted using a Morris Water Maze (Pang et al., 2015). Rats were trained to locate a hidden escape platform (10 x 10cm) located below the water surface in a pool (1.5m diameter). The working memory procedure consisted of 6 trials in one day; each trial had a sample and a choice phase. During the sample phase, rats began from a predetermined location in a zone without the platform and had 60 sec to locate a randomly placed escape platform in the pool. Rats which failed to locate the platform were manually led to the platform. The choice phase commenced 60 seconds after the sample phase and was identical in procedure to the sample phase. Rats spent a minimum of 30 minutes in a holding cage between trials. In each session, the start and the escape platform locations were distributed to equally throughout the pool (3 zones X 2 trials each for a total of 6 trials). Swim paths were recorded for offline analysis of path efficiency (defined as ratio of the straight-line distance between the start and the escape platform location and total distance traveled by a rat; a value of 1 indicates the most efficient path) using ANYmaze software. Rats were tested for spatial working memory performance prior to injury, rats with matched pre-injury path efficiency scores were assigned to either sham or FPI groups such that the average pre-injury path efficiency scores were not different between rats receiving sham or FPI. Rats in each stratum were randomly assigned to an injury group (Sham or FPI) and treatment (saline, LPS-RS*U*, CLI-095) groups. Rats had one session of testing at 1 week (early phase) and 1 month and/or 3 months (late phase) after injury.

### 2.7 Statistical analysis

Statistical analyses were performed using Graphpad Prism 8. A total of 140 rats and 6 mice (pregnant dams) were used in the study. All independent samples were tested for normality and homogeneity of variance in Graphpad Prism 8 using descriptive statistics from Levene’s test. Post-hoc Tukey’s test was used to assess statistical significance of between-group differences. In behavioral studies (Fig. 7 and 8), data from sample and choice phases were analyzed separately using mixed-design ANOVA or two-way repeated measures ANOVA with time as repeated measure variable. Appropriate tests were selected depending upon the samples and their distribution for each experiment and on consultation with the Rutgers Biostatistics core. The significance level was set to *p* < 0.05. Data are shown as mean ± s.e.m. All data and statistical results are presented in Supplemental Tables 1 and 2 respectively.

## 3.0 Results

### 3.1 Differential contribution of glial signaling to TLR4 modulation of excitability in sham and injured rats

The sterile inflammatory response after brain injury is known to enhance TLR4 signaling in the hippocampus (Ahmad et al., 2013; Laird et al., 2014). Brain injury results in dentate hilar neuronal loss and increases in expression of microglia and reactive astrocytes (Gupta et al., 2012; Li et al., 2015; Neuberger et al., 2017a). Our earlier studies identified that TLR4 is expressed in neurons in the hippocampal dentate gyrus and has divergent effects on hippocampal dentate excitability; reducing excitability in controls while enhancing excitability early after brain injury (Li et al., 2015). However, the cellular and molecular signaling mechanisms mediating these divergent effects remain unresolved. To test the contribution of glial signaling to TLR4 modulation of dentate excitability we examined the ability of glial metabolic inhibitors to occlude TLR4 modulation of dentate afferent evoked excitability. As reported previously Li et al. (2015), CLI-095 (also known as TAK-242/Resatorvid, 10ng/ml), a small molecule inhibitor of TLR4 signaling (Matsunaga et al., 2011), increased granule cell population spike amplitude in slices from rats one week after sham injury (Fig. 1A, Note: the CLI095 data was reported in Li et al. (2015) are not shown). In contrast, CLI095 failed to alter granule cell population spike amplitude when the slices were incubated in a drug cocktail containing CLI095 and the metabolic inhibitors (MI) of astrocytes and microglia (1mM fluoroacetate and 50nM minocycline, respectively for 45 min) (Fig 1B, summarized in sham plots in Fig.1E, population spike amplitude in mV in response to a 4mA stimulation, sham in ACSF: 039. ± 0.08, n=8 slices, 3 rats, and sham incubated in CLI-095 and glial MI: 0.28±0.06, n=8 slices, 3 rats, p>0.05 by two-way ANOVA followed by post-hoc Tukey’s test). Thus, glia have a critical role in CLI-095 mediated increase in population spike amplitude in sham controls (Fig. 1F, sham plots).

**Figure 1.**
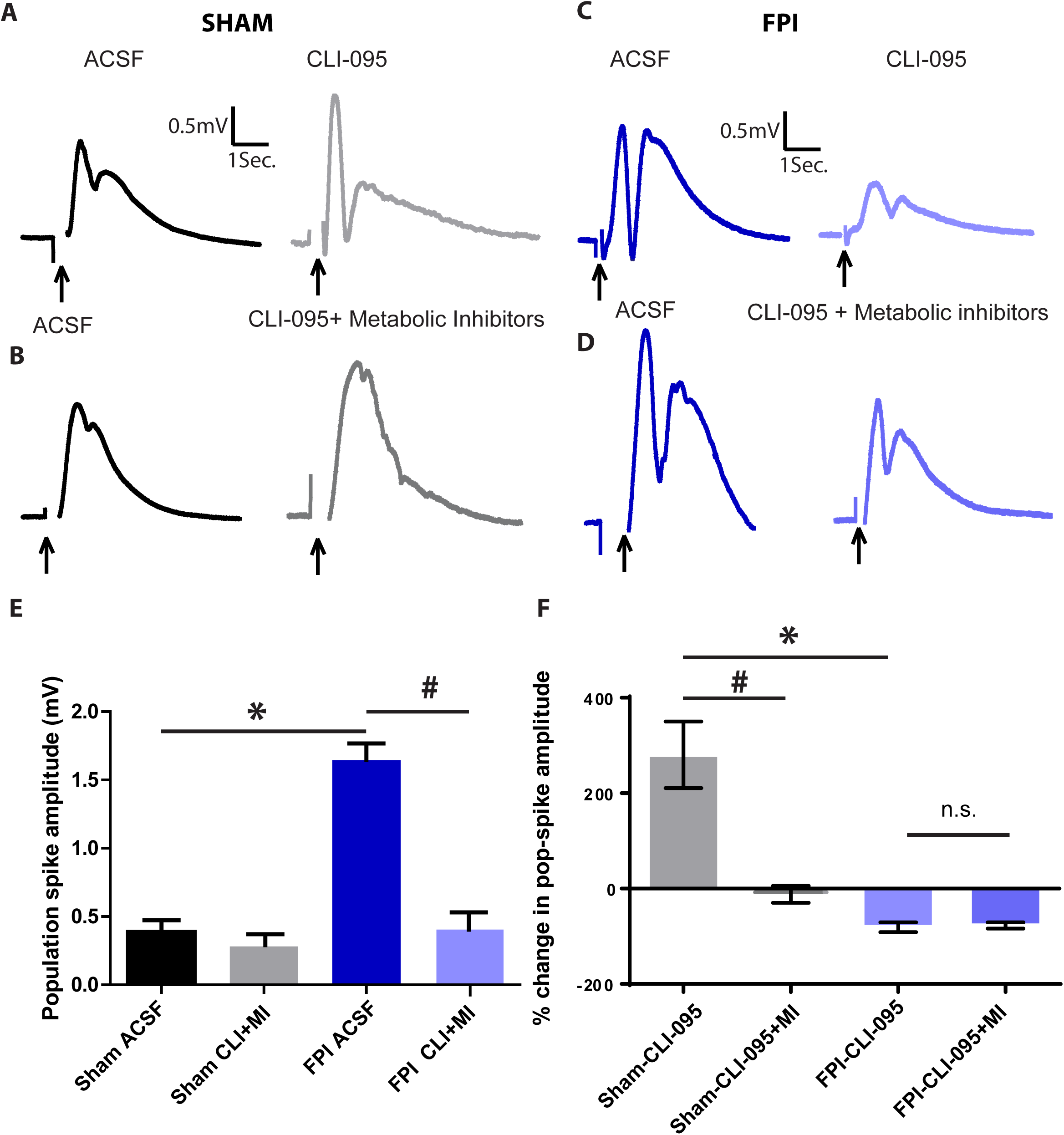
Differential role for glial signaling in TLR4 modulation of dentate excitability in sham and injured rats. (A-D). Granule cell population responses evoked by a 4mA stimulus to the perforant path in slices from sham rats before and after CLI-095 (10ng/ml) in A and before and after incubation in CLI-095+Fluororoacetate (1mM)+Minocycline (50nM) in B. Responses in slices from FPI rats before and after CLI-095 (10ng/ml) in C and before and after incubation in CLI-095+Fluororoacetate (1mM)+Minocycline (50nM) in D. Arrows indicate stimulus artifact. (E-F) Summary plots of population spike amplitude (E) and % change in population spike amplitude compared to corresponding ACSF treatment condition (F). * indicates p<0.05 compared to corresponding recordings in sham, # indicates p<0.05 and n.s. indicates p>0.05 for pairwise comparison with corresponding control drug incubation by TW-ANOVA followed by pairwise Tukey’s test.

One week after brain injury, CLI-095 reduced perforant path-evoked dentate population spike amplitude (Fig. 1C) indicating that TLR4 signaling enhances network excitability after FPI (Li et al., 2015). Unlike findings in sham controls, the reduction in population spike amplitude with CLI-095 treatment persisted unchanged when CLI095 treatment was combined with glial MI (Fig 1D and Fig. 1E FPI data, population spike amplitude in mV in response to a 4mA stimulation, FPI in ACSF: 1.646 ± 0.12, n=8 slices, 3 rats, and FPI incubated in CLI-095 and glial MI: 0.40±0.13, n=8 slices, 3 rats, p>0.05 by two-way ANOVA followed by post-hoc Tukey’s test). These data indicate glial signaling is unlikely to mediate TLR4 enhancement of dentate excitability after brain injury (Fig. 1F, FPI plots). Experiments using a mechanistically distinct TLR4 antagonist, LPS-RS*U* (1µg/ml), confirmed these findings (data not shown). Next we examined whether glial MI could directly impact network excitability. While the population spike amplitude in slices from FPI rats was greater than in sham rats, treatment with glial MI alone did not directly alter network excitability in slices from either sham or FPI rats (Fig. 2A-C, sham in ACSF: 0.29±0.05; sham in glial MI: 0.23± 0.06, -18.33±12.11 % change in n=5 slices, 3 rats; FPI in ACSF: 1.67 ± 0.13, 1.30±3.46% change in n=5 slices, 3 rats; and FPI in glial MI: 1.68±0.12, n=5 slices, 3 rats; p<0.0001 for effect of injury F(1,8)=101.8). Thus, although glial signaling mediates TLR4 modulation of neuronal excitability in sham rats, it does not underlie TLR4 enhancement of excitability after FPI.

**Figure 2.**
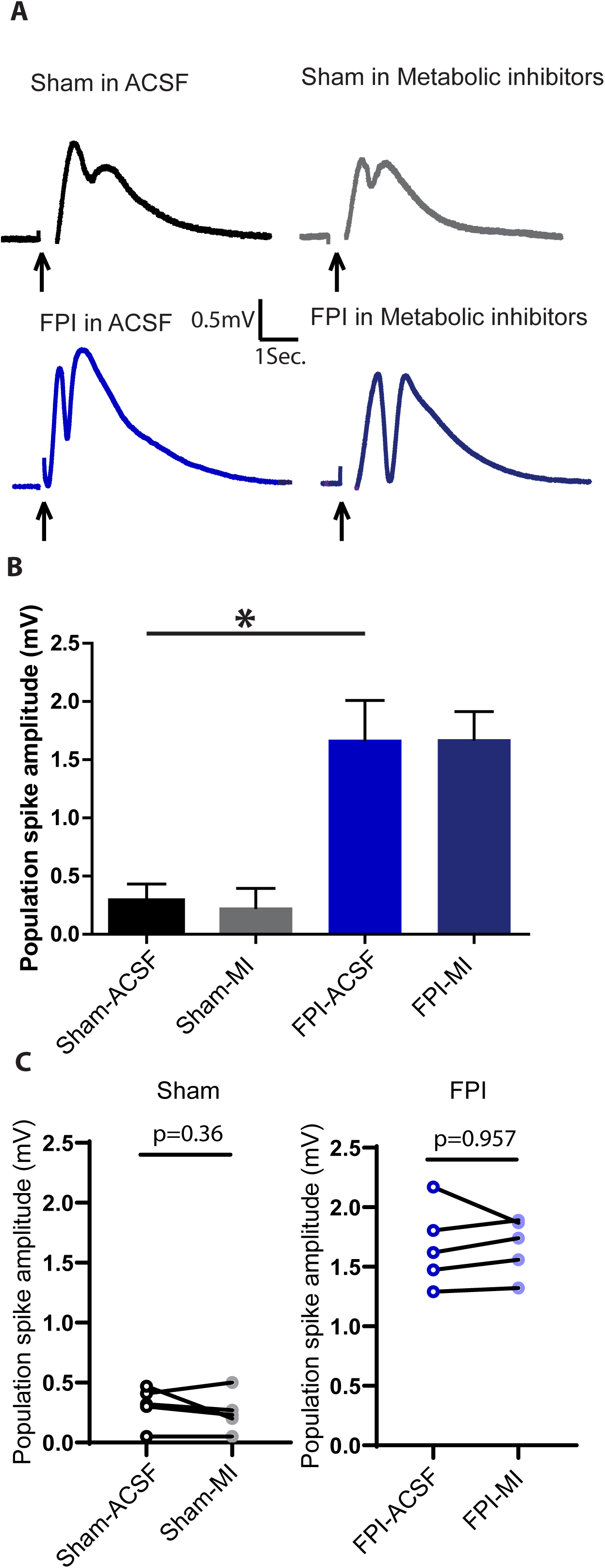
Effect of glial metabolic inhibitors on dentate population spike responses. (A). Granule cell population responses evoked by a 4mA stimulus to the perforant path in slices from the various experimental conditions. Arrows indicate stimulus artifact. (B-C) Summary plots of population spike amplitude (B) and pairwise comparison of population spike amplitude compared to corresponding ACSF treatment condition (C). * indicates p<0.05 by TW-ANOVA followed by pairwise Tukey’s test.

### 3.2 Neuronal signaling underlies TLR4 modulation of AMPAR currents

To directly examine the role for glial versus neuronal signaling in TLR4 modulation of excitability we established pure neuronal cultures from wild-type embryonic mouse hippocampi (Fig. 3A). Cultured cells were confirmed to be purely neuronal by the presence of the microtubule associate protein (MAP2) and absence astrocytic (GFAP) or microglial (Iba1) markers (Fig. 3B). In neuronal cultures, stimulation of neuropil evoked synaptic AMPAR currents, which were enhanced in cultures treated with the TLR4 agonist HMGB1 (10ng/ml) (Fig. 3C-D, AMPAR charge transfer in pA.sec: ACSF: 152.3 ± 6.80, 7 cells from 4 different repeats, HMGB-1: 254.4± 15.57, 7 cells from 4 replicates, p<0.05 by one-way ANOVA followed by a Tukey’s multiple comparison test). The ability of TLR4 agonist to increase excitability is the opposite of what is observed in slices from control rats and similar to observations in *ex vivo* slices from the injured brain (Li et al., 2015). Similarly, when neuronal cultures *in vitro* were treated with the TLR4 antagonist LPS-RS*U* (1µg/ml) the AMPAR current charge transfer was reduced (Fig. 3C-D, AMPAR charge transfer in pA.sec: cultures incubated in LPS-RS*U*: 93.4± 13.75, 6 cells from 4 replicates, p<0.05 versus ACSF by one-way ANOVA followed by a Tukey’s multiple comparison test) as reported in granule cells in *ex vivo* slices from brain injured rats (Li et al., 2015). Once again, these data show that the response of cultured neurons to TLR4 ligands is similar to that observed in granule cells from slices one week after FPI, suggesting that the trauma of dissociation may confer an injury-like phenotype to TLR4 effects in neuronal cultures. The ability of TLR4 ligands to bidirectional modulate of AMPAR currents in the absence of glia, and in a direction similar to that observed in slices after brain injury, is consistent with our finding (Fig. 1) that glial signaling is not necessary for TLR4-dependent enhancement of excitability after brain injury.

**Figure 3.**
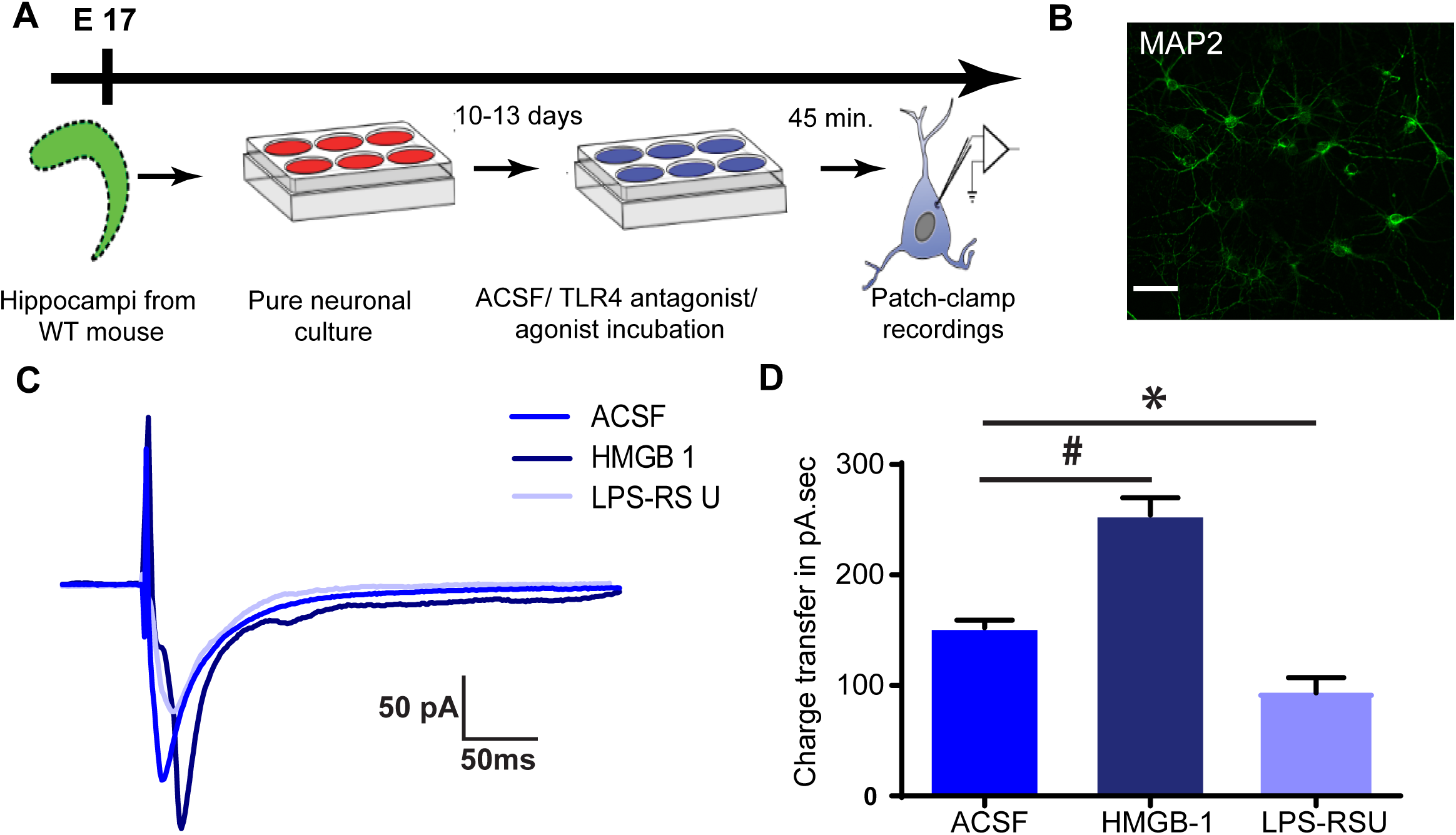
TLR4 modulation of AMPAR currents in hippocampal neurons *in vitro*. (A). Schematic of experimental design shows timeline for preparation of cultures followed by drug treatments and whole cell recordings. (B). Example maximum intensity projection of confocal image stacks from a hippocampal neuronal culture plated at embryonic day 17 and stained for MAP2 at 12 days *in vitro* (DIV) to reveal neurites. (C). Overlay of sample AMPAR current traces recorded in response to neuropil stimulation in neuronal cultures. Neurons were held at -60 mV to obtain recordings in ACSF, TLR4 agonist HMGB1 and TLR4 antagonist LPS-RS*U*. (D). Summary plot of AMPAR current charge transfer. * and # indicates p<0.05 compared to ACSF by One Way ANOVA followed by post-hoc Tukey’s test.

### 3.3 TLR4 antagonist treatment *in vivo* reduces post-injury increases in inflammatory signaling downstream of TLR4

Earlier studies have shown that anti-inflammatory agents improve cell death after brain trauma by suppressing the inflammatory responses mediated downstream of TLR4 signaling (Dong et al., 2011). Our data show that TLR4 signaling reduces excitability in shams and increases excitability in controls. However, whether blocking TLR4 signaling causes opposite changes in downstream signaling pathways in the uninjured and injured brain, which could contribute to the differing effects TLR4 signaling on excitability in sham and FPI rats is not known. To test this possibility, we examined western blots from hippocampal tissue ipsilateral to injury for the endogenous TLR4 ligand, HMGB1, and downstream signaling elements of the MyD88-dependent and independent pathways (See schematic in Supplementary Fig. 2). Fresh hippocampal tissue was obtained three days after injury, from rats that had undergone sham or FPI followed by treatment with saline or CLI-095 treatment (0.5mg/kg, 3 doses at 8-hour intervals starting 20-24 hrs after injury). In saline treated rats, hippocampal expression of HMGB1 was significantly enhanced after FPI compared to sham rats (Fig. 4A-B, p=0.02 by TW ANOVA test, followed by post-hoc pairwise Tukey’s test). While CLI-095 treatment failed to enhance HMGB1 levels in sham controls (p=0.99 for sham-saline vs. sham-CLI), HMGB1 levels in FPI rats treated with CLI-095 trended to decrease when compared to saline-treated FPI rats (p=0.065 for FPI-saline vs. FPI-CLI by pairwise Tukey’s test) and was not different from saline-treated sham rats. Next we examined the pathways downstream of MyD88-dependent TLR4 signaling: MyD88, IkBα, and NFκB (Schematic in Supplementary Fig. 2). Levels of the three proteins downstream of the MyD88-dependent TLR4 pathway were enhanced in saline-treated FPI rats compared to saline-treated sham rats. (Fig. 4C-F). Compared to saline, CLI-095 treatment reduced the level of all three proteins in FPI rats without altering their levels in sham (Fig. 4C-F, Suppl. Table 1-2). Similarly, proteins downstream of the MyD88-independent pathway, TICAM2 and IRF3 were also enhanced in saline-treated FPI rats than in sham controls (Fig. 4G-I; Suppl. Table 1-2). CLI-095 treatment returned TICAM2 and IRF3 levels in FPI rats back to levels comparable to sham controls without altering their levels in sham rats (Fig. 4G-I; Suppl. Table 1-2). Moreover, TNFα, which is downstream of both MyD88-dependent and independent pathways was also enhanced in saline-treated FPI rats and decreased to sham levels after CLI-095 treatment (Fig 4G, J; Supplementary Table 1-2). However, CLI-095 did not alter TNFα levels in sham rats (Fig 4G, J; Supplementary Table 1-2). Thus, *in vivo* CLI-095 treatment significantly reduced hippocampal expression of TLR4 effectors examined in the injured brain while the treatment failed to alter expression of any of the TLR4 effectors tested in sham rats. These findings confirm the ability of *in vivo* treatment with CLI-095 to reduce injury-induced increases in TLR4 signaling in the hippocampus without altering cellular inflammatory signaling in controls.

**Figure 4.**
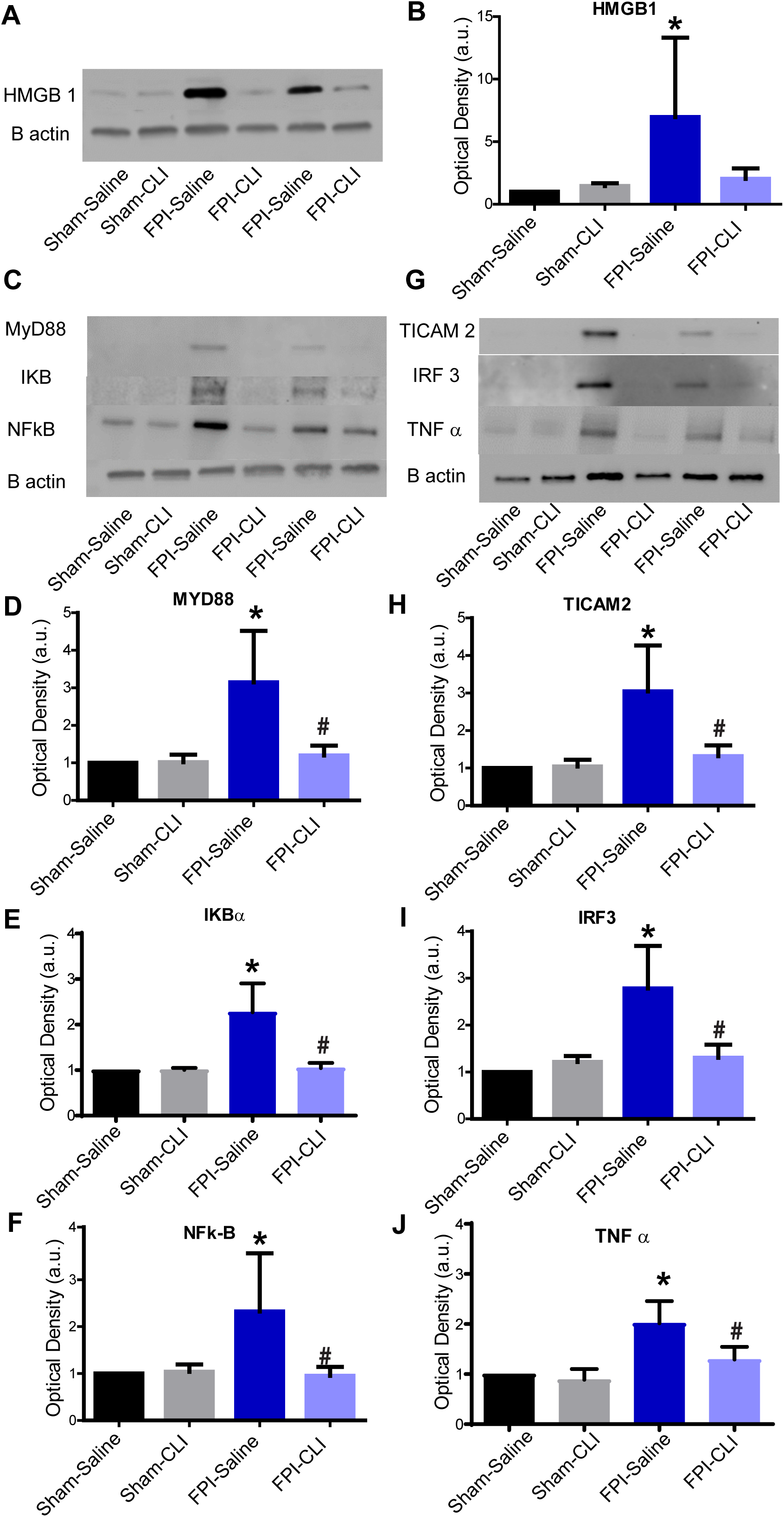
TLR4 antagonism *in vivo* suppresses increases in TLR4 signaling after brain injury. Representative western blots of HMGB1 (A) MyD88, IkBα, and NFκB, (C) and, TICAM2, IRF3 and TNFα (G) in hippocampal samples from the injured side obtained 3 days after vehicle/CLI-095 treatment. Treatments were started 24 hours after injury. Corresponding β-actin bands are illustrated. (A, C and G) Summary histograms of expression of HMGB1 (B), MyD88 (D), IkBα (E), NFκB (F), TICAM2 (H), IRF3 (I) and TNFα (J), normalized to the expression levels in sham-vehicle treated controls. * indicates p<0.05 compared to sham and # indicates p<0.05 compared to corresponding ACSF by TW ANOVA followed by post-hoc Tukey’s test.

### 3.4 Role of TNF-α in TLR4 effects on network excitability

Since TNFα is a direct downstream effector of the MyD88-dependent pathway and is also recruited by MyD88-independent pathways (Akira and Takeda, 2004), we examined whether blocking basal TNFα signaling impacts network excitability in the absence of the TLR4 ligand. In *ex vivo* slices from sham injured rats, incubation in the TNFα antibody (anti-TNFα, 1µg/ml, 1hr) increased dentate population spike amplitude (Fig. 5A-B, population spike amplitude evoked by a 4mA stimulus, in mV, sham: before anti-TNFα 0.02±0.01; after anti-TNFα: 0.33±0.22, n=8 slices from 4 rats, p=0.028 by paired *t*-test). This increase in population spike amplitude in anti-TNFα is similar to what is observed in slices from sham injured rats treated with TLR4 antagonist (Fig. 1A and Li et al., 2015). In contrast to findings in slices from sham rats, incubation in anti-TNFα significantly reduced perforant path-evoked population spike amplitude in slices from FPI rats (Fig. 5C-D, population spike amplitude in mV at 4mA stimulation, FPI before anti-TNFα: 1.43 ± 0.22, FPI after anti-TNFα incubation: 0.422 ± 0.19, n=14 slices from 5 rats, p<0.0001 by paired *t*-test). Blocking TNFα which is elevated levels after brain injury (Fig. 4J and Atkins et al., 2007; Sinha et al., 2017), reduced network excitability, consistent with the effect of TLR4 antagonist application in slices from rats after FPI.

**Figure 5.**
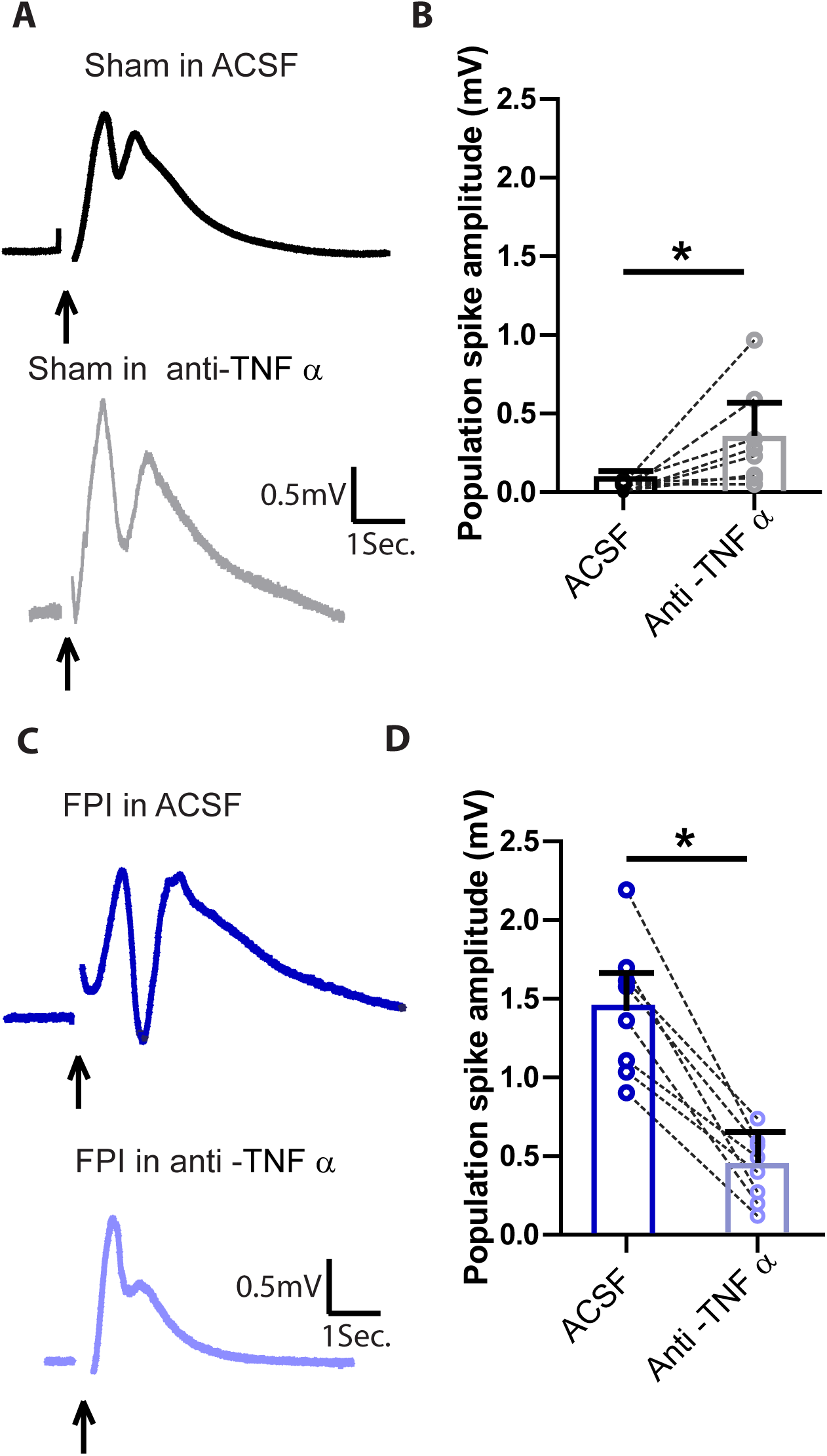
Effect TNFα signaling on network excitability in sham and injured rats. (A) Representative dentate population responses evoked by a 4mA stimulus to the perforant path in a slice from a sham rat before (above) and after (below) incubation in anti-TNFα (1µg/ml). Arrows indicate stimulus artifact. (B) Summary data of effect of anti-TNFα (1µg/ml) on perforant path-evoked granule cell population spike amplitude in slices from sham rats. (C) Example dentate population responses perforant path stimulation at 4 mA in slices from FPI rats before (above) and after (below) incubation in anti-TNFα (1µg/ml). Arrows indicate stimulus artifact. (D) Summary data of effect of anti-TNFα (1µg/ml) on perforant path-evoked granule cell population spike amplitude in slices from sham rats. * indicates p<0.05 by paired Student’s *t*-test.

Since the effects of anti-TNFα on excitability paralleled that of TLR4 antagonists, we examined the potential ability of anti-TNFα to occlude the effects of the TLR4 agonist HMGB1. We previously reported that HMGB1 (10 ng/ml) reduced dentate population spike amplitude in slices from sham rats (Fig 6A-B, Li et al., 2015, HMGB1 data from prior study were used to detrmine % change data in Fig. 6A-B). The ability of HMGB1 to reduce dentate population spike amplitude in slices from sham rats was decreased in the presence of anti-TNFα (1µg/ml) (Fig. 6A-B, % change in population spike amplitude compared to corresponding ACSF, sham-HMGB1: -55.01±9.45 and sham-HMGB1+anti-TNFα: - 27.81±2.59, n=8 slices from 3 rats, p=0.045 by Students *t*-test). In contrast to the increase in dentate population spike amplitude following HMGB1 treatment reported in slices from FPI rats (Fig 6C-D, Li et al., 2015, HMGB1 data from prior study were used to detrmine % change data in Fig. 6C-D), co-treatment with anti-TNFα and HMGB1 decreased dentate population spike amplitude in slices from FPI rats (Fig. 6c, % change in population spike amplitude compared to ACSF, FPI-HMGB1: 77.32±21.59 and FPI-HMGB1+anti-TNFα: -47.51±3.89, n=11 slices from 4 rats, p=0.001 by Students *t*-test). These results suggests that, in spite of the opposite functional effects and differential neuro-glial involvement in TLR4 signaling in sham and FPI rats, TNFα signaling contributes to TLR4 regulation of excitability in both the uninjured brain and after brain injury.

**Figure 6.**
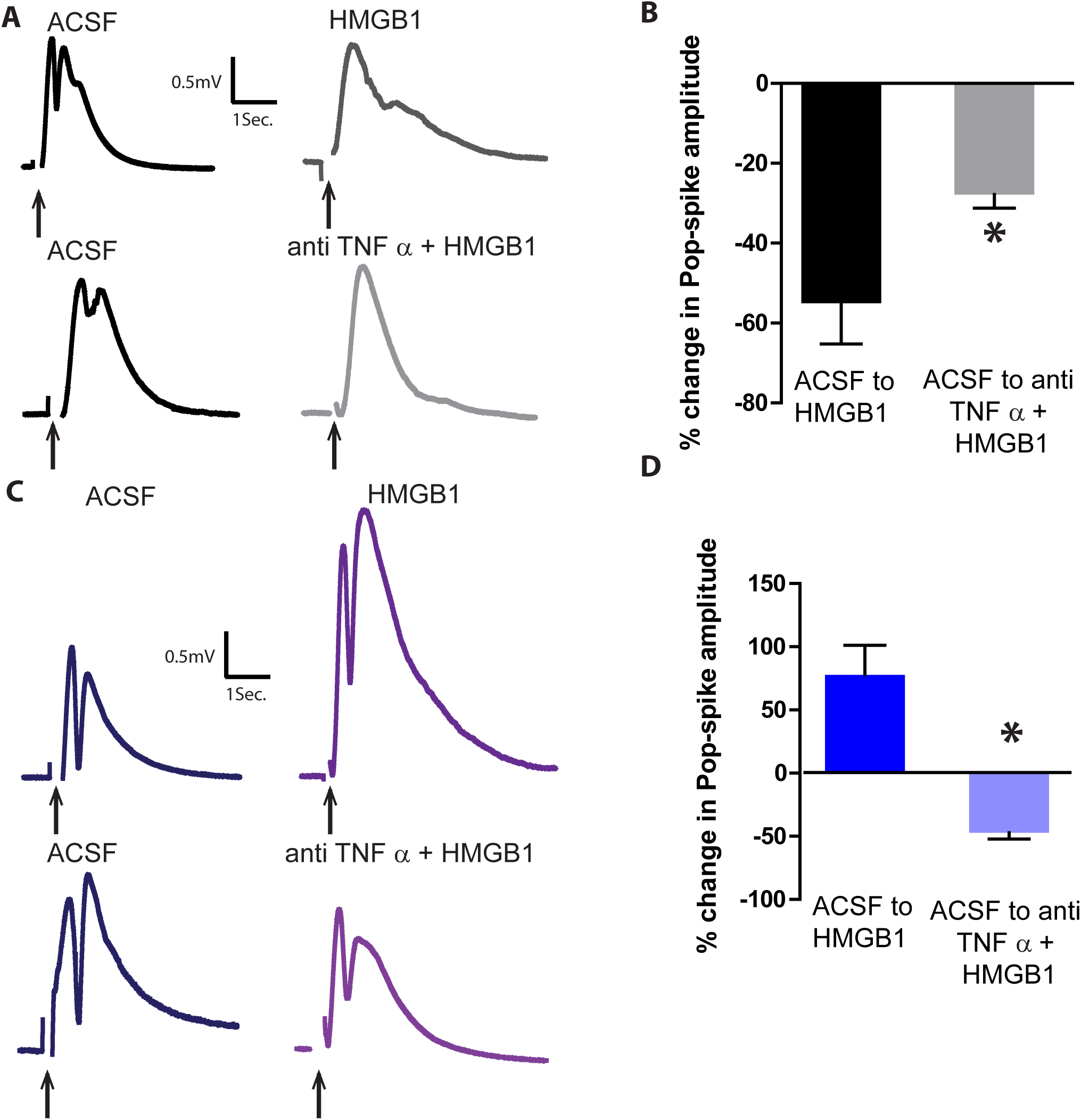
Contribution of TNFα signaling toTLR4 effects on network excitability in sham and injured rats. (A) Representative dentate population responses evoked by a 4mA stimulus to the perforant path in slices from sham rats before and after HMGB1 (10ng/ml) in upper panel and before and after incubation in HMGB1+ anti-TNFα (1µg/ml) in lower panel. Arrows indicate stimulus artifact. (B) Summary of % change in population spike amplitude in the drug(s) compared to corresponding ACSF treatment condition in slices from sham rats 1. (C) Dentate population spike traces evoked by a 4mA stimulus to the perforant path in slices from FPI rats before and after HMGB1 in upper panel and before and after incubation in HMGB1+ anti-TNFα) in lower panel. (D) Summary of % change in population spike amplitude in the drug(s) compared to corresponding ACSF treatment condition in slices from sham rats. Error bars indicate s.e.m. * indicates p<0.05 by unpaired Student’s *t*-test.

**Figure 7.**
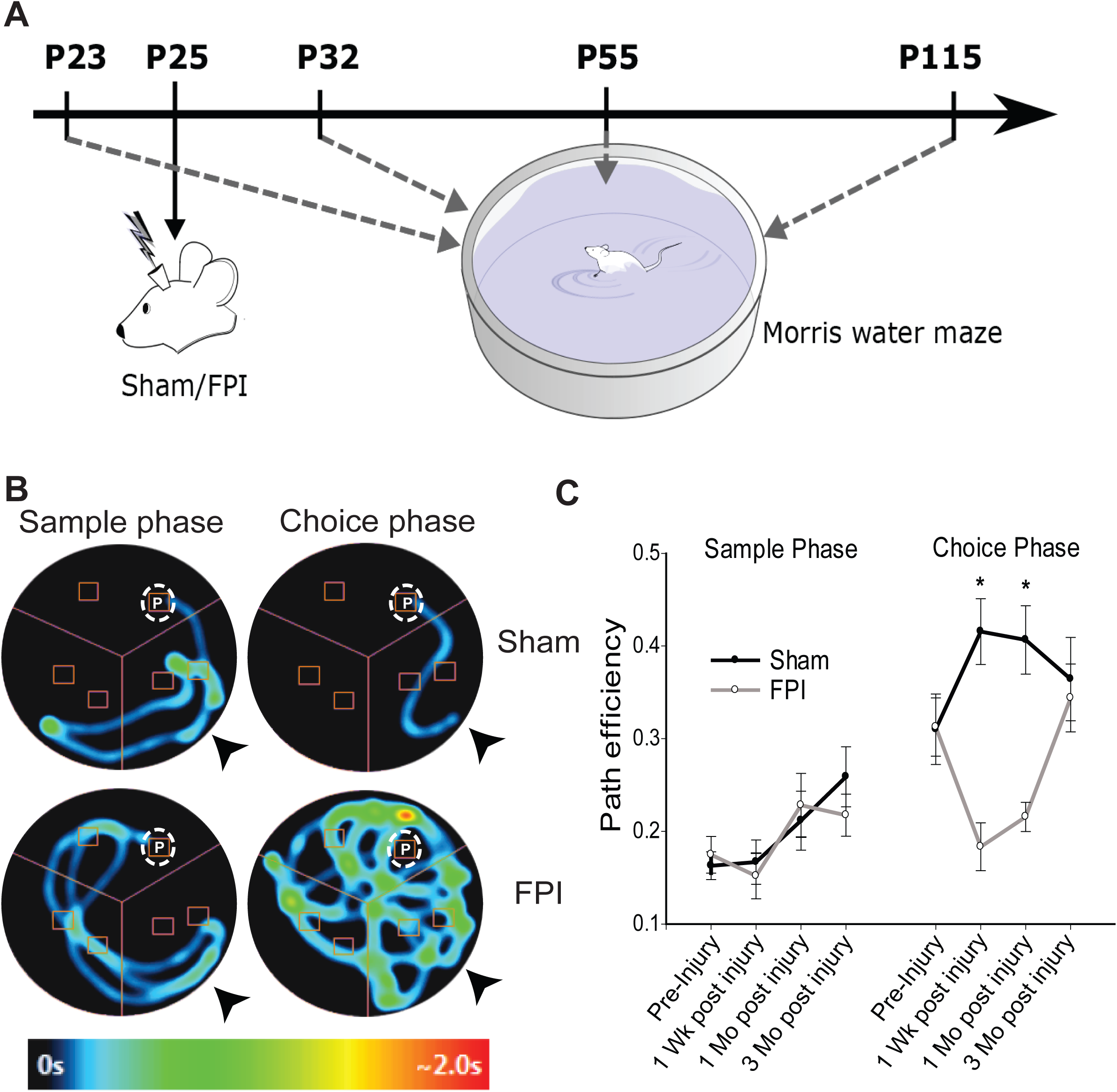
Brain injury impairs working memory function. (A). Schematic of experimental design shows timeline for working memory testing pre and post injury using Morris Water Maze. (B) Representative heatmaps of time spent at each location as the rat traced the path to the platform during sample and choice phase of a delayed match to position task in sham and FPI rats 1 week, after injury. Arrows indicate insertion point in pool. (C) Summary of path efficiency during sample and choice phases, *p<0.05 by mixed design ANOVA followed by pairwise Tukey’s test.

**Figure 8.**
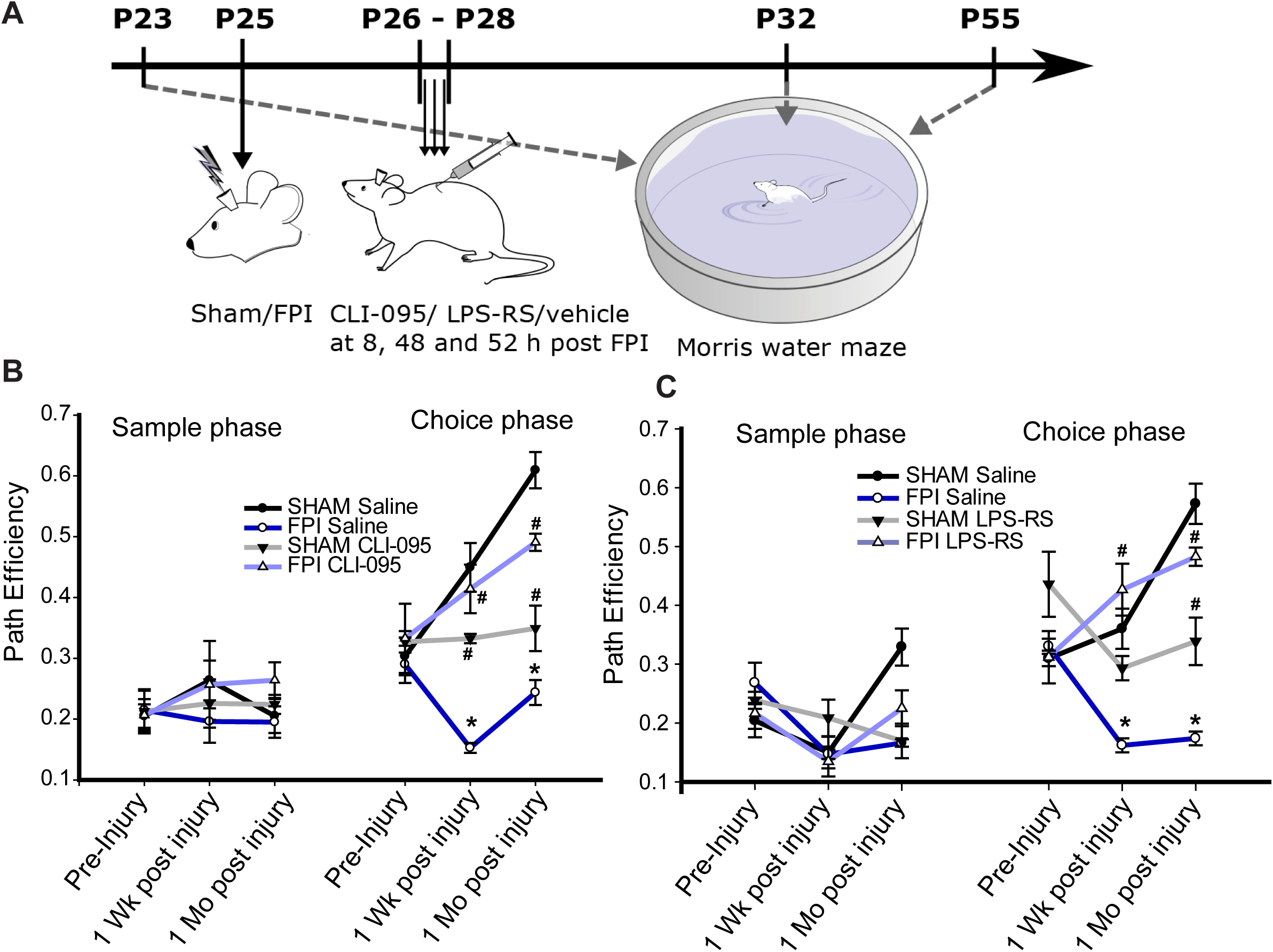
TLR4 antagonist treatment early after FPI improves working memory function. (A) Schematic of experimental design shows timeline for working memory testing pre and post injury using Morris Water Maze. (B) Summary of path efficiency during sample and choice phases of a delayed match to position task in vehicle or CLI-095 treated (systemic) sham and preinjury, 1 week and 1 month after FPI. (C) Summary of path efficiency during sample and choice phases of a delayed match to position task in vehicle or LPS-RS*U* treated (systemic) sham and pre-injury, 1 week and 1 month after FPI. *indicates p<0.05 compared to sham and # indicates p<0.05 compared to corresponding vehicle treatment by TW-RM ANOVA followed by pairwise Tukey’s test, n= 9 rats each group.

### 3.5 TLR4 modulation of network excitability alters working memory function

Since low activity levels in the dentate are crucial for its role in memory processing and TLR4 antagonists lead to opposing changes in dentate excitability in injured and uninjured rats we reasoned that blocking TLR4 signaling would differentially impact memory processing in the uninjured and injured brain. Indeed, brain injury, which is associated with enhanced dentate excitability, is known to affect dentate and hippocampus-dependent memory function (D’Ambrosio et al., 1998; Hamm et al., 1996; Pang et al., 2015; Smith et al., 2015). We focused on a short-term spatial working memory task dependent on dentate function to examine whether FPI, and the associated network alterations, impairs memory processing. Rats were assessed for spatial working memory on a modified Morris Water Maze (MWM) task before injury, at 1 week (Early phase), and at 1 and 3 months (Late phase) after injury. Path efficiency (Fig. 7A, see methods), a measure independent of distances between start and escape platform locations and swimming speed of rats, was used to quantify performance (Pang et al., 2015). To limit confounds, rats with matched pre-injury path efficiency scores were assigned to either sham or FPI groups such that the average pre-injury path efficiency scores were not different between rats receiving sham or FPI. Path efficiency on the sample phase increased as training proceeded, but did not show any significant differences between FPI and sham rats (Fig. 7B-C, path efficiency, sham: pre:0.16±0.01; 1week: 0.166±0.02; 1month: 0.21±0.03; 3month: 0.25±0.03; FPI: pre: 0.17±0.01; 1 week: 0.151±0.02; 1 month: 0.22±0.03; 3 month: 0.21±0.02, n=8 rats each, F_(1,14)_=0.026 and p=0.87 for effect of Injury; F_(3,14)_=3.15 and p=0.057 for effect time and F_(3,14)_=0.215 and p=0.8 for interaction by Mixed design ANOVA). Path efficiency during the choice phase after a 60 sec retention interval was not different between sham and FPI rats prior to injury (Fig. 7C, pre-injury path efficiency, sham: 0.310±0.04, FPI: 0.312±0.03, n=8 rats, p=0.96 by Mixed design ANOVA followed by post hoc Tukey’s test). However, injured rats demonstrated a significant reduction in path efficiency 1 week and 1 month (Fig. 7B-C, path efficiency, sham-1wk: 0.41±0.03; FPI-1wk: 0.183±0.028; sham-1mo: 0.40±0.04; FPI-1mo: 0.21±0.01, n=8 rats/group, p<0.001 for FPI vs. sham at same time point by Mixed design ANOVA followed by post hoc Tukey’s test) which recovered by 3 months (Fig. 7C. Sham-3mo; 0.364±0.05, n=4 rats; FPI-3mo 0.34±0.04, n=4 rats, p>0.99 by Mixed design ANOVA followed by post hoc Tukey’s test). These data identify a transient deficit in memory function 1 week and 1 month after FPI.

Next, we examined whether treatment with a TLR4 antagonist could improve deficits in working memory performance in brain injured rats. Based on the initial findings (Fig 7), we chose to focus on one week and one month after injury to test the effects of pharmacologic treatment (Schematic in Fig. 8A). As before, rats were matched and stratified within the four experimental groups based on pre-injury path efficiency scores. Path efficiency on the sample phase did not show any significant effect of injury/drug or time (Fig. 8B, n=9 rats per group, F_(3,32)_=0.36 and p=0.78 for effect of Injury/Drug; F_(2,32)_=0.3 and p=0.74 for effect time and F_(6,32)_=0.23 and p=0.95 for interaction by TW RM ANOVA). In pre-injury trials, path efficiency during choice phase following a 60 sec retention interval was not different between groups (Fig. 8B, n=9 rats per group, p>0.99 by TW RM ANOVA followed by post-hoc pairwise Tukey’s test). As expected, saline-treated brain injured rats showed an impairment in path efficiency during choice phase compared to sham-saline rats one week and one month after FPI. Consistent with our prediction based on changes in dentate excitability, CLI-095-treatment improved path efficiency in FPI rats compared to saline treated counterparts both one week and one month after injury (Fig. 8B, FPI-saline: 1 week after injury 0.152±0.008 and FPI-CLI-095: 1 week after injury 0.41±0.03 n=9 rats, p<0.001 and FPI-saline: 1 month after injury 0.24±0.020, n=9 rats and FPI-CLI-095 1month after injury 0.49±0.014, n=9 rats, p<0.005 by TW-RM ANOVA followed by pairwise Tukey’s test). In contrast, sham rats treated with systemic CLI-095 showed a reduction in path efficiency compared to sham-saline rats one week and one month after sham-injury/treatment (Fig. 8B sham-saline: 1 week after injury 0.44±0.04 and Sham-CLI-095: 1 week after injury 0.332±0.007 n=9 rats, p=0.04 by post hoc Tukey’s test and sham-saline: 1 month after injury 0.60±0.029, n=9 rats and Sham-CLI-095 1 month after injury 0.349±0.03, n=9 rats, p<0.005 by TW-RM ANOVA followed by pairwise Tukey’s test). While the impairment in memory performance following TLR4 antagonist treatment in uninjured rats is surprising, it is consistent with the ability of CLI-095 to enhance network excitability which would impair dentate memory function.

### 3.6 Focal and transient hippocampal TLR4 antagonism alters working memory function

Systemic administration of CLI-095 can potentially alter central and peripheral immune responses and raises the possibility that the suppression of global immune responses to brain injury may underlie the efficacy of TLR4 modulation on neurobehavioral outcomes after FPI. To limit the potential contribution of systemic effects of TLR4 antagonism, we examined whether local, unilateral hippocampal injection of LPS-RS*U* (5µl of a 2mg/ml solution, intrahippocampal injection 24 hrs after FPI/sham) on the injured side was able to alter working memory performance as observed after systemic treatment. Similar to what was observed after systemic CLI-095 treatment, path efficiency on the sample phase and during pre-injury trials did not show any significant effect of injury/drug or time (Fig. 8C). Focal TLR4 antagonist treatment reduced path efficiency in uninjured controls at one month (Fig. 8C, sham-saline: 1 month after injury 0.57±0.034, n=9 rats and Sham-LPS-RS*U* 1 month after injury 0.338±0.04, n=9 rats, p=0.028 by TW-RM ANOVA followed by pairwise Tukey’s test). Additionally, there was a trend for the treatment to decrease path efficiency one week after sham injury which did not reach statistical significance (Fig. 8C sham-saline: 1 week after injury 0.36±0.033 and Sham-LPS-RS*U*: 1 week after injury 0.29±0.020 n=9 rats, p=0.74 by TW-RM ANOVA followed by pairwise Tukey’s test). In contrast, LPS-RS*U*-treatment improved path efficiency in brain injured FPI rats compared to saline treated counterparts both one week and one month after injury (Fig. 8C, FPI-saline: 1week after injury 0.162±0.008 and FPI-LPS-RS*U*: 1week after injury 0.426±0.04 n=9 rats, p<0.001 and FPI-saline: 1 month after injury 0.173±0.011, n=9 rats and FPI-LPS-RS*U* 1month after injury 0.482±0.015, n=9 rats, p<0.005 by TW-RM ANOVA followed by pairwise Tukey’s test). Thus, as with systemic CLI-095, focal LPS-RS*U* improved memory performance one week and one month after FPI while impairing path efficiency at the one-month time point in sham controls (Fig. 8C). Together, these data support our hypothesis that early changes in dentate excitability and TLR4 modulation of dentate excitability contribute to neurological deficits including impaired working memory performance after brain injury.

## 4.0 Discussion

### 4.1 Janus Faced effects of TLR4 signaling on excitability and memory function

This study identifies that distinct cellular mediators contribute to the divergent effects of TLR4 on excitability in the normal and injured brain which have a lasting impact on behavioral outcomes. The data show that the ‘Janus faced’ effects of TLR4 on excitability are determined by differential recruitment of glial versus neuronal TLR4 signaling.

It has long been recognized that sparse firing is an essential functional feature of the dentate gyrus and its ability to ‘gate’ activity is critical for memory processing and for limiting pathology (Acsady and Kali, 2007; Dengler and Coulter, 2016; Lothman et al., 1992). TLR4 activation enhances dentate network excitability after brain injury which would compromise the dentate gate. Consequently, TLR4 antagonists reduce dentate excitability after brain injury and improve working memory function *in vivo*. Simultaneously, TLR4 antagonist treatment *in vivo* reduces injury-induced increases in molecular signals associated with both the MyD88-dependent and MyD88-independant pathways effectively suppressing TLR4-dependent immune response to injury. Following brain injury, TLR4 antagonists reduced dentate excitability in *ex vivo* slices even in the presence of glial metabolic inhibitors. Moreover, AMPAR currents in neuronal cultures depleted of glia were enhanced by HMGB1, a TLR4 agonist, and reduced by LPS-RS*U*, a selective TLR4 antagonist, demonstrating a role for neuronal TLR4 signaling in augmenting excitability in the injured brain. Interestingly, blocking TNFα, a proinflammatory cytokine downstream of MyD88-dependent TLR4 signaling, reduced dentate excitability and occluded the ability of HMGB1 to increase excitability indicating that neuronal TLR4 signaling acts through TNFα to enhance excitability after brain injury.

In contrast, TLR4 signaling appears to reinforce the dentate gate in the uninjured brain, where TLR4 signaling limits dentate network excitability during afferent stimulation (Li et al., 2015). Consistently, blocking basal TLR4 signaling enhances dentate excitability in response to input activation in *ex vivo* slices and compromises working memory function *in vivo*, demonstrating that basal TLR4 signaling regulates memory processing. Distinct from the effects on excitability, *in vivo* treatment with TLR4 antagonist failed to alter effectors downstream of the MyD88-dependent and MyD88-indepandant pathways in controls, indicating that antagonists did not have paradoxical effects on inflammatory signaling. Unlike after injury, glial metabolic inhibitors blocked TLR4 antagonist enhancement of dentate excitability in slices from sham rats indicating involvement of glial signaling in TLR4 modulation of excitability in controls. Blocking TNFα tended to enhance excitability suggesting a role for basal TNFα signaling in limiting network excitability. Anti-TNFα reduced the ability of a TLR4 agonist to decrease excitability indicating that TNFα is downstream of TLR4 signaling in controls. These findings coupled with the ability of glial MI to block effects of TLR4 modulation in sham rats suggest that TLR4 recruitment of signaling pathways in glia and glia-derived TNFα contribute to suppression of basal excitability in the uninjured dentate gyrus. Thus, the Janus faces of TLR4 signaling in the brain are distinguished by recruitment of glial metabolic processes and TNFα to limit excitability and support working memory function in the uninjured brain while activation of neuronal TLR4 and TNFα in the injured brain increases network excitability and compromises working memory function.

### 4.2 Immune signaling and memory function

Several lines of evidence point to multiple roles for molecules, once thought to be confined to the peripheral immune system, in regulating normal function in healthy brains and in neurological disorders (Pickering and O’Connor, 2007; Williamson and Bilbo, 2013). Basal levels of neuroimmune signaling through specific cytokines such as TNFα have been shown to contribute to synaptic plasticity and memory processing (Avital et al., 2003; Balschun et al., 2004; Ben Menachem-Zidon et al., 2011; Goshen et al., 2007). Studies in transgenic mice lacking specific immune receptors, including TLR4, have identified memory and cognitive changes underscoring the importance of neuroimmune interactions in shaping brain function (Okun et al., 2012; Potter et al., 2019). Since neuroimmune interplay is critical for the developmental establishment of circuits (Szepesi et al., 2018), use of germline knockouts confounds the ability to distinguish between immune regulation of circuit development and basal neuroimmune signaling in behaviors. Here we show that even a brief and transient block of basal TLR4 signaling, which increases network excitability, leads to prolonged impairments in working memory function for up to a month after treatment. Moreover, two mechanistically distinct TLR4 antagonists administered either focally in the hippocampus (LPS-RS*U*) or systemically (CLI-095) had similar effects in impairing working memory function, reducing the potential for off target effects. Curiously, an earlier study in mice reported impaired memory performance one week after a single treatment with the exogenous TLR4 agonist LPS and that TLR4 antagonist treatment abolished LPS induced memory impairments and suppressed the associated inflammatory response (Zhang et al., 2018). These findings contrast with what we would expect based on results demonstrating that TLR4 antagonist impairs memory function in uninjured rats (Fig. 8). However, it should be noted that LPS treatment in Zhang et al. (2018) enhanced inflammatory cytokines and TLR4 mediated signaling and may better reflect the neuroimmune environment present in the injured brain in our study.

We find that TLR4 antagonist treatment reduced network excitability and improved working memory function after brain injury. The behavioral outcomes paralleled the opposing effect of TLR4 ligands on excitability rather than the effects on inflammatory signaling which were unchanged in controls. Thus, it is reasonable to propose that the changes in excitability drive underlie the effects on memory function. Indeed, in controls, TLR4 antagonists enhanced excitability which would be expected to compromise sparse neuronal activity and impair working memory performance (Madar et al., 2019; Scharfman and Bernstein, 2015; Wixted et al., 2014). In contrast, when neuronal excitability is increased following brain injury, TLR4 antagonists reduced excitability, potentially aiding in maintaining sparse neuronal activity, which improved working memory. These data suggest that there may be an optimal range for TLR4 signaling to maintain physiological levels of excitability; both blocking basal levels, as with TLR4 antagonism in controls, as well as enhanced TLR4 signaling, as occurs after injury, can be pathological. Our findings are in line with the ability of innate immune signals to regulate the immediate early gene Arc and reports that Arc expression both above and below the optimal range can impair synaptic plasticity and cognitive function (Pickering and O’Connor, 2007; Rosi, 2011). Curiously, anti-TNFα mimics the effects TLR4 antagonists in slices from both the control and injured brains. Moreover, TLR4 mediated decrease in excitability in controls and increase in excitability after FPI appear to involve TNFα signaling suggesting that there may be a physiological range for TNFα that regulates excitability and synaptic plasticity at optimal levels. Our findings are consistent with the ability of exogenous TNFα to modulate synaptic excitability and plasticity (Belarbi et al., 2012; Maggio and Vlachos, 2018; Stellwagen et al., 2005). However, HMGB1 effects on dentate excitability in controls were reduced but not eliminated in anti-TNFα. It is possible HMGB1 may recruit additional receptor and or downstream pathways to impact network excitability.

Traumatic brain injury is known to impair memory and cognitive function (Azouvi et al., 2017; Hamm et al., 1996; Lyeth et al., 1990; Titus et al., 2016; Vallat-Azouvi et al., 2007). In earlier studies, mild lateral FPI (1.2 atm) was shown to contribute to transient working memory deficits which recovered by 3 weeks (Pang et al., 2015). At the moderate injury strength (2-2.2 atm) used in the current study working memory deficits persist 4 weeks after injury and recover by 3 months, which is consistent with increasing neuropathology with injury severity. There is considerable evidence for the ability of non-specific immune modulators to limit post-traumatic inflammatory responses including TLR4 expression and improve neurological outcomes (Mao et al., 2012; Wang et al., 2015; Yu et al., 2012). Our study using both local and systemic administration of specific TLR4 antagonists clearly demonstrates that suppression of TLR4 is beneficial following experimental brain injury in rats. The paradoxical detrimental effects of TLR4 antagonist treatment in the uninjured brain, interestingly, are coupled to an increase in network excitability by mechanisms that are currently unknown. However, our data indicate that TLR4 does not modulate AMPA and NMDA receptor currents in slices from uninjured rats (Li et al., 2015), suggesting that modulation of GABA currents or intrinsic neuronal excitability need to be examined in future studies. Regardless of the underlying mechanisms, the effects of TLR4 antagonists on excitability and memory function in uninjured brain warrants caution while proposing to use TLR4 antagonists for therapeutics.

### 4.2 Post-injury switch in cellular mediators of TLR4 effects on excitability

We previously identified that post-traumatic increase in TLR4 expression is predominantly neuronal (Li et al., 2015). However, activation of glial purinergic signaling by TLR4 is known to acutely increase excitatory synaptic drive to CA1 neurons by modulating metabotropic glutamate receptors (Pascual et al., 2012). Additionally, LPS induced neurotoxicity in cortical cultures was found to be mediated through glia (Lehnardt et al., 2003). Moreover, glial TLR4 signaling through endogenous TLR4 ligands such as HMGB1, can enhance proinflammatory cytokines such as IL-1β and TNFα in immune cells including microglia and promote excitability (Maroso et al., 2010). Our study using two complementary methods: glial metabolic inhibitors and isolated neuronal cultures, demonstrates that TLR4 modulation of AMPAR currents is independent of glia. It is possible that the lack of TLR4 in cortical neurons (Lehnardt et al., 2003) or use of LPS, which activates a broader immune response, contributes to the glial dependence of TLR4 signaling in prior studies. It is notable that our data in control slices demonstrate that glial signaling is necessary for maintaining basal levels of excitability and TLR4 modulation of excitability in the uninjured brain.

Our findings suggest two distinct cellular and signaling pathways for TLR4 function depending on whether there is injury to the brain. Classically, TLR4 signaling in glia is mediated by MyD88-TNFα dependent pathway and MyD88-independent pathway. TLR4 activation induces a MyD88-dependent increase in TNFα (Gao et al., 2009) and is proposed to enhance NMDA currents (Balosso et al., 2014; Maroso et al., 2010) and recruit glial purinergic receptors (Pascual et al., 2012). However, the post-traumatic increase in dentate excitability is mediated by enhancement of polysynaptic AMPAR currents with no change in NMDAR currents (Santhakumar et al., 2000). Consistent with this data, TLR4 enhances granule cell AMPAR and not NMDAR currents after FPI (Li et al., 2015). Interestingly, glial TNFα has been shown to recruit Calcium permeable AMPARs (CP-AMPARs) to the synapse (Pickering et al., 2005; Pribiag and Stellwagen, 2014; Stellwagen and Malenka, 2006). Since our results with glial metabolic inhibitors and neuronal cultures indicate that TLR4-mediated increases in excitability after injury does not involve glial signaling, it is possible that TLR4 activation after brain injury recruits “neuronal” TNF-α signaling. This would constitute a novel function for neuronal TNFα. TNFα is a likely candidate for the neuropathological effects of TLR4 after brain injury since it is known to contribute to excitotoxicity (Pickering et al., 2005; Stellwagen, 2011). Indeed, an inhibitor of TNFα synthesis has been shown to reverse cognitive deficits induced by chronic neuroinflammation (Belarbi et al., 2012). Unlike the neuronal TLR4 signaling identified after injury, our data suggest that classical glial-TNFα signaling may reduce excitability in the uninjured hippocampus by mechanisms of which are currently under investigation.

### 4.3 Basal TLR4 modulation of network function and behaviors

A particularly interesting finding is the ability of transient TLR4 antagonism to increase excitability and impair working memory performance in control, uninjured animals. These data suggest a critical neurophysiological function for the low levels of TLR4 signaling observed in the uninjured brain. Moreover, the lack of change in immune signaling alongside changes in excitability observed after TLR4 antagonism in the controls suggests that non-immune responses underlie this effect. Yet, blocking glial signaling eliminated the ability of CLI-095 to increase excitability in controls suggesting that glia are necessary intermediaries for this process. We demonstrate changes in memory function following acute and transient modulation of TLR4. These results extend prior works demonstrating a developmental role for TLR4 in learning and memory function in mice with constitutive TLR4 knockout (Okun et al., 2012). It remains to be seen whether TLR4 can modulate glial glutamate uptake and glutamate-glutamine cycling in the brain as has been shown in spinal circuits during LPS challenge (Yan et al., 2015). Regardless, the basal role for TLR4 in modulating dentate excitability and its impact on memory function need to be considered while proposing TLR4 modulators for use in therapies (Gnjatic et al., 2010; Hanke and Kielian, 2011).

## 5.0 Conclusions

Our data identify that TLR4 signaling in neurons contributes to pathological increase in excitability after brain injury by enhancing AMPAR currents through a mechanism that involves TNFα. Blocking TLR4 signaling early after brain injury reduces injury-induced decreases in hippocampal working memory function. However, we find that blocking basal TLR4 signaling in the uninjured brain augments dentate excitability through recruitment of glial signaling and TLR4 demonstrating a role for basal TLR4 signaling in maintaining sparse dentate activity. Thus, blocking the basal TLR4 signaling impairs working memory function. Taken together, the potential therapeutic effect of TLR4 antagonists may be better harnessed by isolating and selectively targeting the distinct pathological signaling downstream of neuronal TLR4.

## 6.0 Acknowledgements

We thank Drs. Roman Shirakov and Stella Elkabes for help with culture studies. We thank Drs. Kelly Hamilton, Deepak Subramainian and Ms. Susan Nguyen for discussions.

---

**Supplemental Figure 1.**
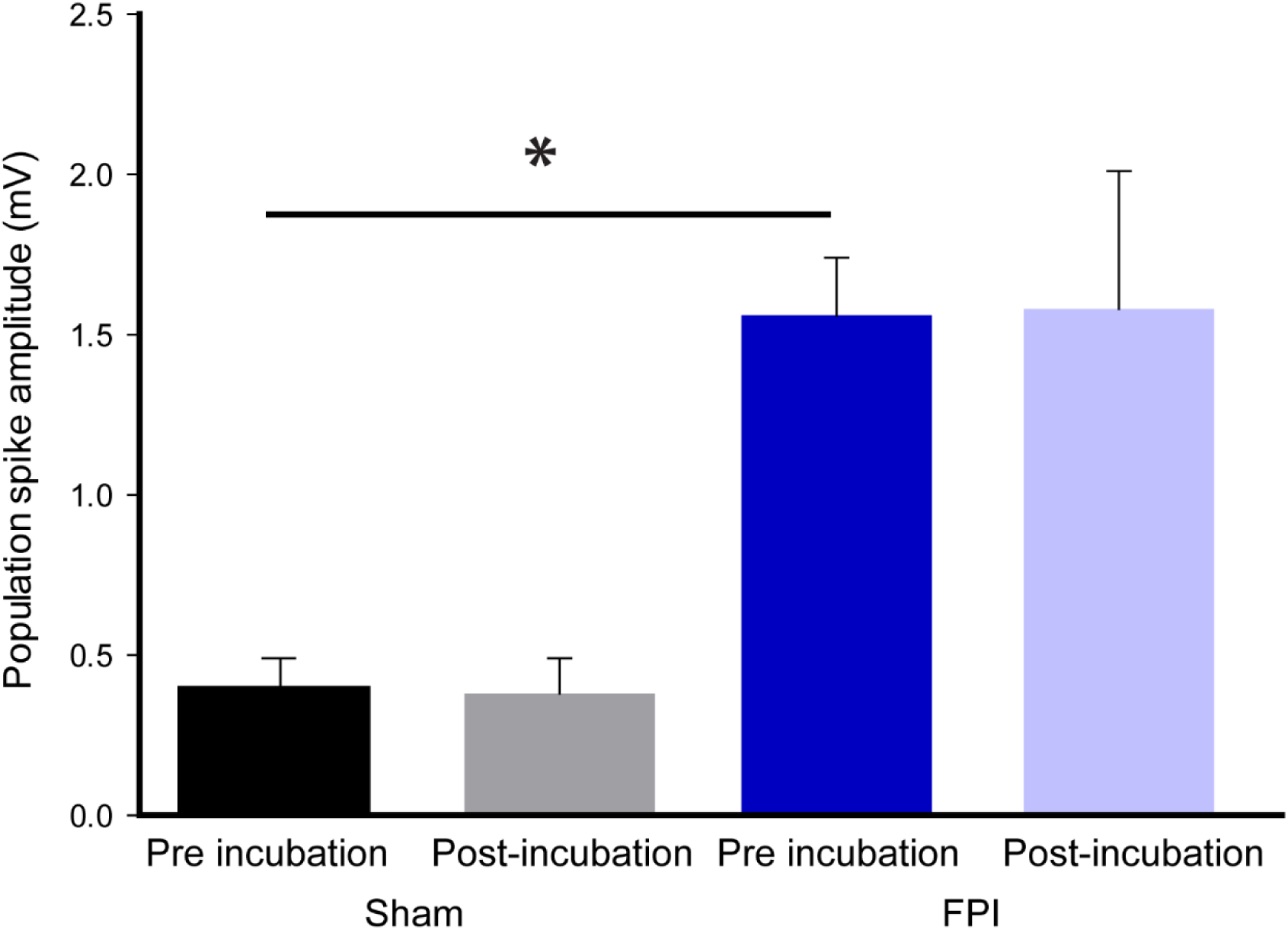
Control data show that vehicle incubations do not alter population spike amplitudes. A). Summary plots of population spike amplitude responses evoked by a 4-mA stimulus to the perforant path in slices from the various experimental conditions. * and # indicates p<0.05.

**Supplemental Figure 2.**
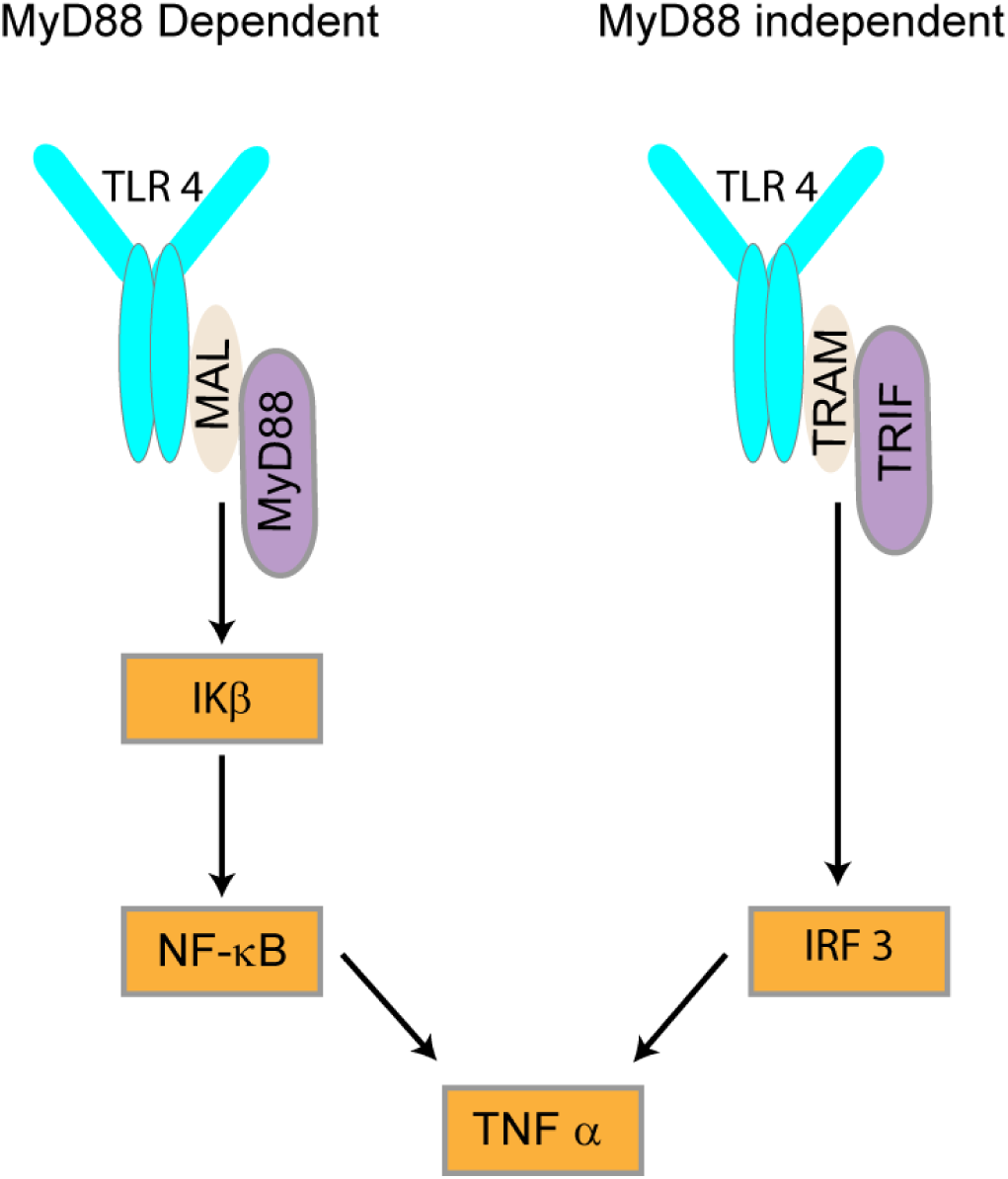
Schematic of MyD88-dependent and MyD88 independent TLR4 pathway. Simplified schematic illustrates the molecular pathways involved in TLR4 signaling. MyD88: Myeloid Differentiation factor 88; MAL: MyD88 adapter like; IkBα: inhibitor of NF-κB; NF-κB: nuclear factor-kappa B; TNFα: Tumor Necrosis Factor α; TRIF: TIR-domain-containing adapter-inducing interferon-β; TICAM2/TRAM: TRIF-related adaptor molecule (TRAM, also known as TIR domain-containing adapter molecule 2 or TICAM2); IRF3: Interferon regulatory factor 3. Adapted from Akira and Takeda (2004)

**Supplemental Table 1.**
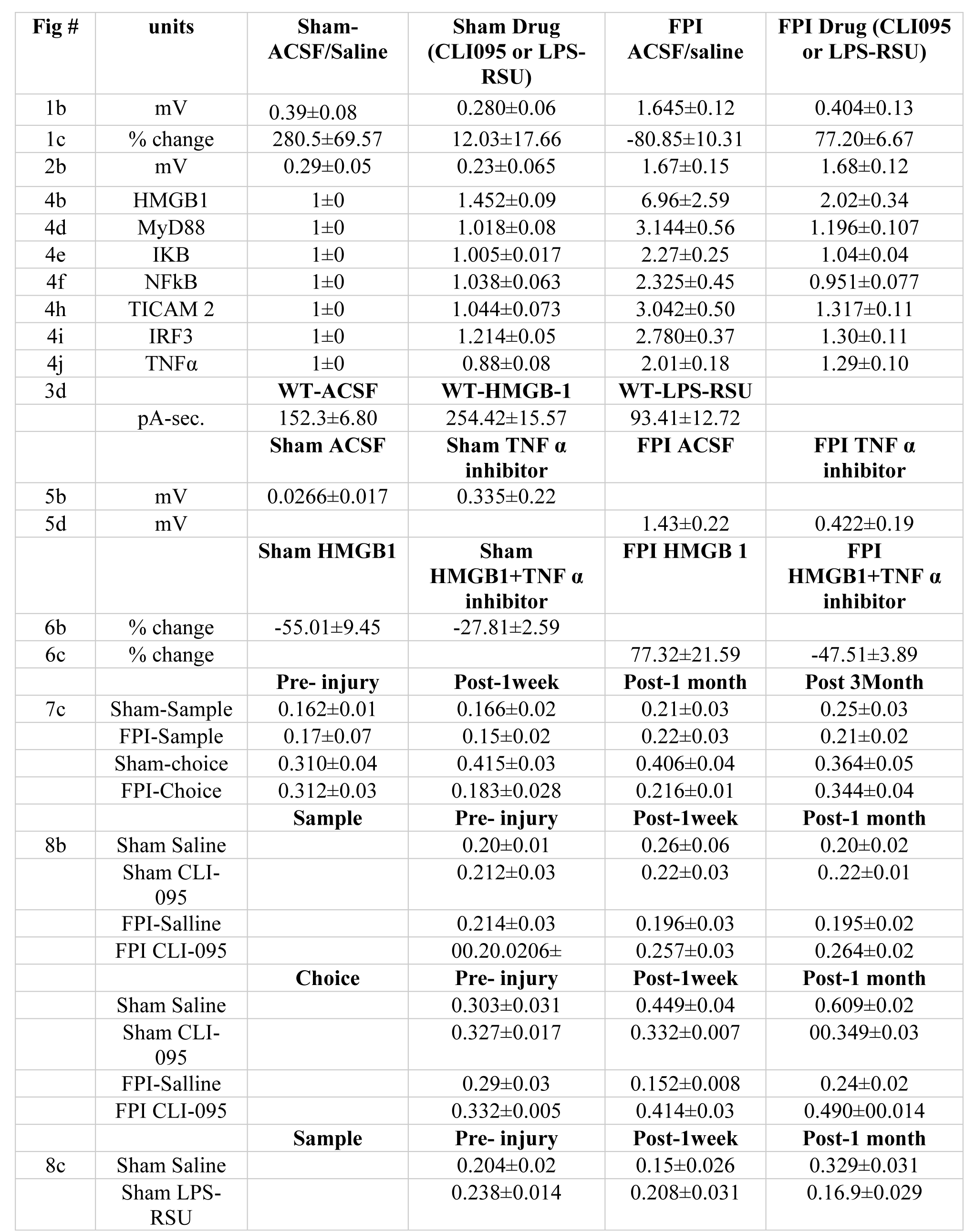
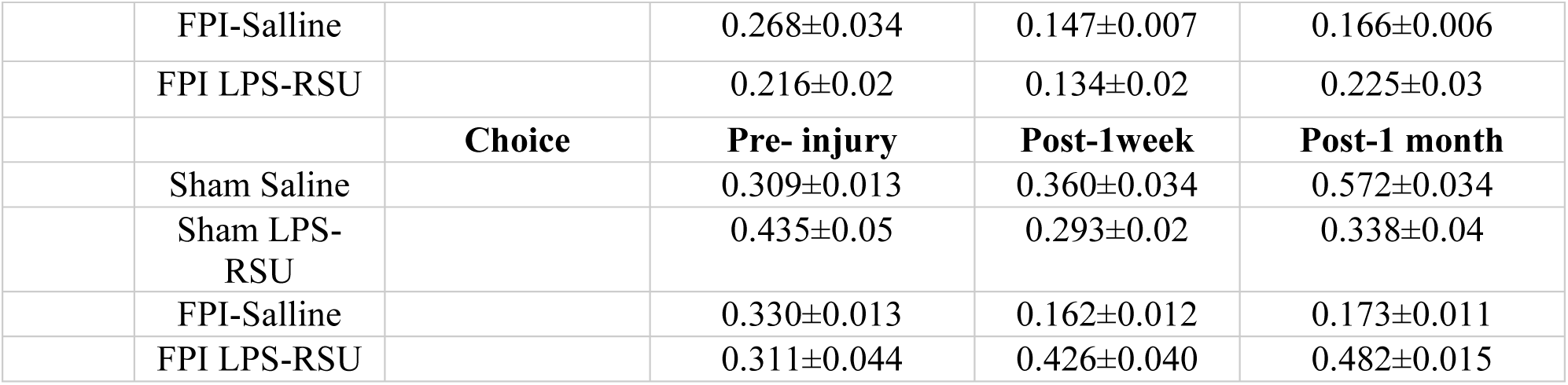
Summary Table of Data

**Supplemental Table 2.** Summary Table of Statistics

|  | Test used | n |  | P Value, DEGREES OF FREEDOM & F/t/z/R/ETC VALUE |  |  |
| --- | --- | --- | --- | --- | --- | --- |
| Figure numbe | Which test | Exact value | Defined | Exact Value |  |  |
| 1b | Two way RM ANOVA test, followed by a Tukey's multiple comparison test | 8,8 | No. of Slices from at least 3 rats/group | Source of Variation<br>Interaction<br>Injury<br>Drug | P value<br>P<0.0001<br>P<0.0001<br>P<0.0001 | F (DFn, DFd)<br>F (1, 14) = 64.04<br>F (1, 14) = 30.64<br>F (1, 14) = 91.88 |
| 1 c | Two way ANOVA test, followed by a Tukey's multiple comparison test | 5, 8,6,9 | No. of Slices from at least 3 rats/group | Source of Variation<br>Interaction<br>Injury<br>Drug | P value<br>P<0.0001<br>P<0.0001<br>P<0.0001 | F (DFn, DFd)<br>F (1, 24) = 28.80<br>F (1, 24) = 60.72<br>F (1, 24) = 27.93 |
| 2b | Two way RM ANOVA test, followed by a Tukey's multiple comparison test | 5,5 | No. of Slices from at least 3 rats/group | Source of Variation<br>Interaction<br>Injury<br>Drug | P value<br>P=0.5069<br>P<0.0001<br>P=0.5653 | F (DFn, DFd)<br>F (1, 8) = 0.482<br>F (1, 8) = 101.8<br>F (1, 8) = 0.359 |
| 2 c | Paired T- Test | 5,5 | No. of Slices from at least 3 rats/group | P=0.36 | T=1.011 |  |
| 2 d | Paired T- Test | 5,5 | No. of Slices from at least 3 rats/group | P=0.957 | T=0.056 |  |
| 3 d | One way ANOVA test, followed by a Tukey's multiple comparison test | 7,7,6 | No. of cells from at least 4 different well plates per group | P<0.0001 | F(2,17)=41.66 |  |
| 4b | TW ANOVA test, followed by a Tukey's multiple comparison test | 6 in each group | No. of rats per group | Source of Variation<br>Interaction<br>Injury<br>Drug | P value<br>P=0.0528<br>P=0.0214<br>P=0.1022 | F (DFn, DFd)<br>F(1,20)=4.23<br>F(1,20)=6.23<br>F(1,20)=2.93 |
| 4d | TW ANOVA test, followed by a Tukey's multiple comparison test | 6 in each group | No. of rats per group | Source of Variation<br>Interaction<br>Injury<br>Drug | P value<br>P=0.0028<br>P=0.007<br>P=0.0032 | F (DFn, DFd)<br>F(1,20)=11.65<br>F(1,20)=16.24<br>F(1,20)=11.22 |
| 4e | TW ANOVA test, followed by a Tukey's multiple comparison test | 6 in each group | No. of rats per group | Source of Variation<br>Interaction<br>Injury | P value<br>P=0.0001<br>P<0.0001 | F (DFn, DFd)<br>F(1,20)=21.80<br>F(1,20)=24.67 |
|  |  |  |  | Drug | P=0.0002 | F(1,20)=21.42 |
| 4f | TW ANOVA test,<br>followed by a Tukey's<br>multiple comparison test | 6 in<br>each<br>group | No. of rats<br>per group | **Source of**<br>**Variation**<br>Interaction<br>Injury<br>Drug | **P value**<br>P=0.0067<br>P=0.0152<br>P=0.0096 | **F (DFn, DFd)**<br>F(1,20)=9.147<br>F(1,20)=7.049<br>F(1,20)=8.188 |
| 4h | TW ANOVA test,<br>followed by a Tukey's<br>multiple comparison test | 6 in<br>each<br>group | No. of rats<br>per group | **Source of**<br>**Variation**<br>Interaction<br>Injury<br>Drug | **P value**<br>P=0.0028<br>P=0.0002<br>P=0.0041 | **F (DFn, DFd)**<br>F(1,20)=11.63<br>F(1,20)=19.92<br>F(1,20)=10.49 |
| 4i | TW ANOVA test,<br>followed by a Tukey's<br>multiple comparison test | 6 in<br>each<br>group | No. of rats<br>per group | **Source of**<br>**Variation**<br>Interaction<br>Injury<br>Drug | **P value**<br>P=0.0003<br>P=0.0001<br>P=0.0042 | **F (DFn, DFd)**<br>F(1,20)=18.70<br>F(1,20)=22.86<br>F(1,20)=10.43 |
| 4j | TW ANOVA test,<br>followed by a Tukey's<br>multiple comparison test | 6 in<br>each<br>group | No. of rats<br>per group | **Source of**<br>**Variation**<br>Interaction<br>Injury<br>Drug | **P value**<br>P=0.0132<br>P<0.0001<br>P=0.0014 | **F (DFn, DFd)**<br>F(1,20)=7.39<br>F(1,20)=39.28<br>F(1,20)=13.66 |
| 5 b | Paired T- Test | 5,5 | No. of Slices<br>from at least<br>3 rats/group | P=0.0283 | T=2.75 |  |
| 5 d | Paired T- Test | 5,5 | No. of Slices<br>from at least<br>3 rats/group | P<0.0001 | T=7.94 |  |
| 6 b | Student's T- Test | 6,5 | No. of Slices<br>from at least<br>3 rats/group | P=0.045 | T=2.322 |  |
| 6 d | Student's T- Test | 6,5 | No. of Slices<br>from at least<br>3 rats/group | P=0.001 | T=4.76 |  |
| 7 c | Mixed design ANOVA<br>test, followed by a<br>Tukey's multiple<br>comparison test | 8,8 | No. of rats<br>per group | **Source of**<br>**Variation**<br>Interaction<br>Injury-drug<br>Time point | **P value**<br>P=0.0063<br>P=0.0004<br>P=0.6368 | **F (DFn, DFd)**<br>F(3,14)=7.39<br>F(1,14)=39.28<br>F(3,14)=13.66 |
| 8 b | Two way RM ANOVA<br>test, followed by a<br>Tukey's multiple<br>comparison test | 9 in<br>each<br>group | No. of rats<br>per group | **Source of**<br>**Variation**<br>Interaction<br>Injury-drug<br>Time point | **P value**<br>P=0.0016<br>P=0.0013<br>P<0.0001 | **F (DFn, DFd)**<br>F(6,32)=4.548<br>F(3,32)=16.56<br>F(2,32)=7.991 |
| 8 c | Two way RM ANOVA<br>test, followed by a<br>Tukey's multiple<br>comparison test | 9 in<br>each<br>group | No. of rats<br>per group | **Source of**<br>**Variation**<br>Interaction<br>Injury-drug<br>Time point | **P value**<br>P<0.0001<br>P<0.0001<br>P=0.0196 | **F (DFn, DFd)**<br>F(6,32)=6.532<br>F(3,32)=12.45<br>F(2,32)=4.346 |

## References

Acsady, L., Kali, S., 2007. Models, structure, function: the transformation of cortical signals in the dentate gyrus. Prog Brain Res 163, 577–599.

Ahmad, A., Crupi, R., Campolo, M., Genovese, T., Esposito, E., Cuzzocrea, S., 2013. Absence of TLR4 reduces neurovascular unit and secondary inflammatory process after traumatic brain injury in mice. PLoS One 8, e57208.

Akira, S., Takeda, K., 2004. Toll-like receptor signalling. Nat Rev Immunol 4, 499–511.

Atkins, C.M., Oliva, A.A., Jr., Alonso, O.F., Pearse, D.D., Bramlett, H.M., Dietrich, W.D., 2007. Modulation of the cAMP signaling pathway after traumatic brain injury. Exp Neurol 208, 145–158.

Avital, A., Goshen, I., Kamsler, A., Segal, M., Iverfeldt, K., Richter-Levin, G., Yirmiya, R., 2003. Impaired interleukin-1 signaling is associated with deficits in hippocampal memory processes and neural plasticity. Hippocampus 13, 826–834.

Azouvi, P., Arnould, A., Dromer, E., Vallat-Azouvi, C., 2017. Neuropsychology of traumatic brain injury: An expert overview. Rev Neurol (Paris) 173, 461–472.

Balosso, S., Liu, J., Bianchi, M.E., Vezzani, A., 2014. Disulfide-Containing High Mobility Group Box-1 Promotes N-Methyl-d-Aspartate Receptor Function and Excitotoxicity by Activating Toll-Like Receptor 4-Dependent Signaling in Hippocampal Neurons. Antioxidants & redox signaling.

Balschun, D., Wetzel, W., Del Rey, A., Pitossi, F., Schneider, H., Zuschratter, W., Besedovsky, H.O., 2004. Interleukin-6: a cytokine to forget. FASEB J 18, 1788–1790.

Beattie, E.C., Stellwagen, D., Morishita, W., Bresnahan, J.C., Ha, B.K., Von Zastrow, M., Beattie, M.S., Malenka, R.C., 2002. Control of synaptic strength by glial TNFalpha. Science 295, 2282–2285.

Belarbi, K., Jopson, T., Tweedie, D., Arellano, C., Luo, W., Greig, N.H., Rosi, S., 2012. TNF-alpha protein synthesis inhibitor restores neuronal function and reverses cognitive deficits induced by chronic neuroinflammation. J Neuroinflammation 9, 23.

Ben Menachem-Zidon, O., Avital, A., Ben-Menahem, Y., Goshen, I., Kreisel, T., Shmueli, E.M., Segal, M., Ben Hur, T., Yirmiya, R., 2011. Astrocytes support hippocampal-dependent memory and long-term potentiation via interleukin-1 signaling. Brain Behav Immun 25, 1008–1016.

Chiu, C.C., Liao, Y.E., Yang, L.Y., Wang, J.Y., Tweedie, D., Karnati, H.K., Greig, N.H., Wang, J.Y., 2016. Neuroinflammation in animal models of traumatic brain injury. J Neurosci Methods 272, 38–49.

Costello, D.A., Watson, M.B., Cowley, T.R., Murphy, N., Murphy Royal, C., Garlanda, C., Lynch, M.A., 2011. Interleukin-1alpha and HMGB1 mediate hippocampal dysfunction in SIGIRR-deficient mice. The Journal of neuroscience : the official journal of the Society for Neuroscience 31, 3871–3879.

D’Ambrosio, R., Maris, D.O., Grady, M.S., Winn, H.R., Janigro, D., 1998. Selective loss of hippocampal long-term potentiation, but not depression, following fluid percussion injury. Brain Res 786, 64–79.

Dengler, C.G., Coulter, D.A., 2016. Normal and epilepsy-associated pathologic function of the dentate gyrus. Prog Brain Res 226, 155–178.

Dong, X.Q., Yu, W.H., Hu, Y.Y., Zhang, Z.Y., Huang, M., 2011. Oxymatrine reduces neuronal cell apoptosis by inhibiting Toll-like receptor 4/nuclear factor kappa-B-dependent inflammatory responses in traumatic rat brain injury. Inflammation research : official journal of the European Histamine Research Society … [et al.] 60, 533–539.

Gao, Y., Fang, X., Tong, Y., Liu, Y., Zhang, B., 2009. TLR4-mediated MyD88-dependent signaling pathway is activated by cerebral ischemia-reperfusion in cortex in mice. Biomed Pharmacother 63, 442–450.

Gnjatic, S., Sawhney, N.B., Bhardwaj, N., 2010. Toll-like receptor agonists: are they good adjuvants? Cancer J 16, 382–391.

Goshen, I., Kreisel, T., Ounallah-Saad, H., Renbaum, P., Zalzstein, Y., Ben-Hur, T., Levy-Lahad, E., Yirmiya, R., 2007. A dual role for interleukin-1 in hippocampal-dependent memory processes. Psychoneuroendocrinology 32, 1106–1115.

Gupta, A., Elgammal, F.S., Proddutur, A., Shah, S., Santhakumar, V., 2012. Decrease in tonic inhibition contributes to increase in dentate semilunar granule cell excitability after brain injury. J Neurosci 32, 2523–2537.

Hamm, R.J., Temple, M.D., Pike, B.R., O’Dell, D.M., Buck, D.L., Lyeth, B.G., 1996. Working memory deficits following traumatic brain injury in the rat. Journal of neurotrauma 13, 317–323.

Hanke, M.L., Kielian, T., 2011. Toll-like receptors in health and disease in the brain: mechanisms and therapeutic potential. Clin Sci (Lond) 121, 367–387.

Jeltsch, H., Bertrand, F., Lazarus, C., Cassel, J.C., 2001. Cognitive performances and locomotor activity following dentate granule cell damage in rats: role of lesion extent and type of memory tested. Neurobiol Learn Mem 76, 81–105.

Kawai, T., Akira, S., 2007. TLR signaling. Semin Immunol 19, 24–32.

Kielian, T., 2006. Toll-like receptors in central nervous system glial inflammation and homeostasis. J.Neurosci.Res. 83, 711–730.

Laird, M.D., Shields, J.S., Sukumari-Ramesh, S., Kimbler, D.E., Fessler, R.D., Shakir, B., Youssef, P., Yanasak, N., Vender, J.R., Dhandapani, K.M., 2014. High mobility group box protein-1 promotes cerebral edema after traumatic brain injury via activation of toll-like receptor 4. Glia 62, 26–38.

Lehnardt, S., Massillon, L., Follett, P., Jensen, F.E., Ratan, R., Rosenberg, P.A., Volpe, J.J., Vartanian, T., 2003. Activation of innate immunity in the CNS triggers neurodegeneration through a Toll-like receptor 4-dependent pathway. Proc Natl Acad Sci U S A 100, 8514–8519.

Li, Y., Korgaonkar, A.A., Swietek, B., Wang, J., Elgammal, F.S., Elkabes, S., Santhakumar, V., 2015. Toll-like receptor 4 enhancement of non-NMDA synaptic currents increases dentate excitability after brain injury. Neurobiol Dis 74, 240–253.

Lothman, E.W., Stringer, J.L., Bertram, E.H., 1992. The dentate gyrus as a control point for seizures in the hippocampus and beyond. Epilepsy Res Suppl 7, 301–313.

Lyeth, B.G., Jenkins, L.W., Hamm, R.J., Dixon, C.E., Phillips, L.L., Clifton, G.L., Young, H.F., Hayes, R.L., 1990. Prolonged memory impairment in the absence of hippocampal cell death following traumatic brain injury in the rat. Brain Res 526, 249–258.

Madar, A.D., Ewell, L.A., Jones, M.V., 2019. Temporal pattern separation in hippocampal neurons through multiplexed neural codes. PLoS Comput Biol 15, e1006932.

Maggio, N., Vlachos, A., 2018. Tumor necrosis factor (TNF) modulates synaptic plasticity in a concentration-dependent manner through intracellular calcium stores. J Mol Med (Berl) 96, 1039–1047.

Mao, S.S., Hua, R., Zhao, X.P., Qin, X., Sun, Z.Q., Zhang, Y., Wu, Y.Q., Jia, M.X., Cao, J.L., Zhang, Y.M., 2012. Exogenous administration of PACAP alleviates traumatic brain injury in rats through a mechanism involving the TLR4/MyD88/NF-kappaB pathway. Journal of neurotrauma 29, 1941–1959.

Maroso, M., Balosso, S., Ravizza, T., Liu, J., Aronica, E., Iyer, A.M., Rossetti, C., Molteni, M., Casalgrandi, M., Manfredi, A.A., Bianchi, M.E., Vezzani, A., 2010. Toll-like receptor 4 and high-mobility group box-1 are involved in ictogenesis and can be targeted to reduce seizures. Nat.Med. 16, 413–419.

Matsunaga, N., Tsuchimori, N., Matsumoto, T., Ii, M., 2011. TAK-242 (resatorvid), a small-molecule inhibitor of Toll-like receptor (TLR) 4 signaling, binds selectively to TLR4 and interferes with interactions between TLR4 and its adaptor molecules. Mol Pharmacol 79, 34–41.

Neuberger, E.J., Abdul-Wahab, R., Jayakumar, A., Pfister, B.J., Santhakumar, V., 2014. Distinct effect of impact rise times on immediate and early neuropathology after brain injury in juvenile rats. Journal of Neuroscience Research.

Neuberger, E.J., Gupta, A., Subramanian, D., Korgaonkar, A.A., Santhakumar, V., 2017a. Converging early responses to brain injury pave the road to epileptogenesis. J Neurosci Res.

Neuberger, E.J., Swietek, B., Corrubia, L., Prasanna, A., Santhakumar, V., 2017b. Enhanced Dentate Neurogenesis after Brain Injury Undermines Long-Term Neurogenic Potential and Promotes Seizure Susceptibility. Stem Cell Reports 9, 972–984.

Okun, E., Barak, B., Saada-Madar, R., Rothman, S.M., Griffioen, K.J., Roberts, N., Castro, K., Mughal, M.R., Pita, M.A., Stranahan, A.M., Arumugam, T.V., Mattson, M.P., 2012. Evidence for a developmental role for TLR4 in learning and memory. PLoS One 7, e47522.

Okun, E., Griffioen, K.J., Mattson, M.P., 2011. Toll-like receptor signaling in neural plasticity and disease. Trends Neurosci 34, 269–281.

Pang, K.C., Sinha, S., Avcu, P., Roland, J.J., Nadpara, N., Pfister, B., Long, M., Santhakumar, V., Servatius, R.J., 2015. Long-lasting suppression of acoustic startle response after mild traumatic brain injury. Journal of neurotrauma 32, 801–810.

Pascual, O., Ben Achour, S., Rostaing, P., Triller, A., Bessis, A., 2012. Microglia activation triggers astrocyte-mediated modulation of excitatory neurotransmission. Proc Natl Acad Sci U S A 109, E197–205.

Pickering, M., Cumiskey, D., O’Connor, J.J., 2005. Actions of TNF-alpha on glutamatergic synaptic transmission in the central nervous system. Exp Physiol 90, 663–670.

Pickering, M., O’Connor, J.J., 2007. Pro-inflammatory cytokines and their effects in the dentate gyrus. Prog Brain Res 163, 339–354.

Potter, O.V., Giedraitis, M.E., Johnson, C.D., Cox, M.N., Kohman, R.A., 2019. Young and aged TLR4 deficient mice show sex-dependent enhancements in spatial memory and alterations in interleukin-1 related genes. Brain Behav Immun 76, 37–47.

Pribiag, H., Stellwagen, D., 2014. Neuroimmune regulation of homeostatic synaptic plasticity. Neuropharmacology 78, 13–22.

Roof, R.L., Hall, E.D., 2000. Gender differences in acute CNS trauma and stroke: neuroprotective effects of estrogen and progesterone. Journal of neurotrauma 17, 367–388.

Rosi, S., 2011. Neuroinflammation and the plasticity-related immediate-early gene Arc. Brain Behav Immun 25 Suppl 1, S39–49.

Saletti, P.G., Ali, I., Casillas-Espinosa, P.M., Semple, B.D., Lisgaras, C.P., Moshe, S.L., Galanopoulou, A.S., 2019. In search of antiepileptogenic treatments for post-traumatic epilepsy. Neurobiol Dis 123, 86–99.

Santhakumar, V., Bender, R., Frotscher, M., Ross, S.T., Hollrigel, G.S., Toth, Z., Soltesz, I., 2000. Granule cell hyperexcitability in the early post-traumatic rat dentate gyrus: the ’irritable mossy cell’ hypothesis. J Physiol 524 Pt 1, 117–134.

Santhakumar, V., Ratzliff, A.D., Jeng, J., Toth, K., Soltesz, I., 2001. Long-term hyperexcitability in the hippocampus after experimental head trauma. Ann.Neurol. 50, 708–717.

Scharfman, H.E., Bernstein, H.L., 2015. Potential implications of a monosynaptic pathway from mossy cells to adult-born granule cells of the dentate gyrus. Front Syst Neurosci 9, 112.

Semple, B.D., Zamani, A., Rayner, G., Shultz, S.R., Jones, N.C., 2018. Affective, neurocognitive and psychosocial disorders associated with traumatic brain injury and post-traumatic epilepsy. Neurobiol Dis.

Sinha, S.P., Avcu, P., Spiegler, K.M., Komaravolu, S., Kim, K., Cominski, T., Servatius, R.J., Pang, K.C.H., 2017. Startle suppression after mild traumatic brain injury is associated with an increase in pro-inflammatory cytokines, reactive gliosis and neuronal loss in the caudal pontine reticular nucleus. Brain Behav Immun 61, 353–364.

Smith, C.J., Xiong, G., Elkind, J.A., Putnam, B., Cohen, A.S., 2015. Brain Injury Impairs Working Memory and Prefrontal Circuit Function. Front Neurol 6, 240.

Stellwagen, D., 2011. The contribution of TNFalpha to synaptic plasticity and nervous system function. Adv Exp Med Biol 691, 541–557.

Stellwagen, D., Beattie, E.C., Seo, J.Y., Malenka, R.C., 2005. Differential regulation of AMPA receptor and GABA receptor trafficking by tumor necrosis factor-alpha. J Neurosci 25, 3219–3228.

Stellwagen, D., Malenka, R.C., 2006. Synaptic scaling mediated by glial TNF-alpha. Nature 440, 1054–1059.

Szepesi, Z., Manouchehrian, O., Bachiller, S., Deierborg, T., 2018. Bidirectional Microglia-Neuron Communication in Health and Disease. Front Cell Neurosci 12, 323.

Titus, D.J., Wilson, N.M., Freund, J.E., Carballosa, M.M., Sikah, K.E., Furones, C., Dietrich, W.D., Gurney, M.E., Atkins, C.M., 2016. Chronic Cognitive Dysfunction after Traumatic Brain Injury Is Improved with a Phosphodiesterase 4B Inhibitor. J Neurosci 36, 7095–7108.

Vallat-Azouvi, C., Weber, T., Legrand, L., Azouvi, P., 2007. Working memory after severe traumatic brain injury. J Int Neuropsychol Soc 13, 770–780.

van Vliet, E.A., Aronica, E., Vezzani, A., Ravizza, T., 2018. Review: Neuroinflammatory pathways as treatment targets and biomarker candidates in epilepsy: emerging evidence from preclinical and clinical studies. Neuropathol Appl Neurobiol 44, 91–111.

Vezzani, A., French, J., Bartfai, T., Baram, T.Z., 2011. The role of inflammation in epilepsy. Nat Rev Neurol 7, 31–40.

Wang, C.X., Xie, G.B., Zhou, C.H., Zhang, X.S., Li, T., Xu, J.G., Li, N., Ding, K., Hang, C.H., Shi, J.X., Zhou, M.L., 2015. Baincalein alleviates early brain injury after experimental subarachnoid hemorrhage in rats: possible involvement of TLR4/NF-kappaB-mediated inflammatory pathway. Brain Res 1594, 245–255.

Williamson, L.L., Bilbo, S.D., 2013. Chemokines and the hippocampus: a new perspective on hippocampal plasticity and vulnerability. Brain Behav Immun 30, 186–194.

Wixted, J.T., Squire, L.R., Jang, Y., Papesh, M.H., Goldinger, S.D., Kuhn, J.R., Smith, K.A., Treiman, D.M., Steinmetz, P.N., 2014. Sparse and distributed coding of episodic memory in neurons of the human hippocampus. Proc Natl Acad Sci U S A 111, 9621–9626.

Xavier, G.F., Oliveira-Filho, F.J., Santos, A.M., 1999. Dentate gyrus-selective colchicine lesion and disruption of performance in spatial tasks: difficulties in “place strategy” because of a lack of flexibility in the use of environmental cues? Hippocampus 9, 668–681.

Yan, X., Jiang, E., Weng, H.R., 2015. Activation of toll like receptor 4 attenuates GABA synthesis and postsynaptic GABA receptor activities in the spinal dorsal horn via releasing interleukin-1 beta. J Neuroinflammation 12, 222.

Ye, Y., Xu, H., Zhang, X., Li, Z., Jia, Y., He, X., Huang, J.H., 2014. Association between toll-like receptor 4 expression and neural stem cell proliferation in the hippocampus following traumatic brain injury in mice. Int J Mol Sci 15, 12651–12664.

Yu, J., Swietek, B., Proddutur, A., Santhakumar, V., 2016. Dentate cannabinoid-sensitive interneurons undergo unique and selective strengthening of mutual synaptic inhibition in experimental epilepsy. Neurobiol Dis 89, 23–35.

Yu, W.H., Dong, X.Q., Hu, Y.Y., Huang, M., Zhang, Z.Y., 2012. Ginkgolide B reduces neuronal cell apoptosis in the traumatic rat brain: possible involvement of toll-like receptor 4 and nuclear factor kappa B pathway. Phytother Res 26, 1838–1844.

Zhang, J., Yu, C., Zhang, X., Chen, H., Dong, J., Lu, W., Song, Z., Zhou, W., 2018. Porphyromonas gingivalis lipopolysaccharide induces cognitive dysfunction, mediated by neuronal inflammation via activation of the TLR4 signaling pathway in C57BL/6 mice. J Neuroinflammation 15, 37.

Zhu, H.T., Bian, C., Yuan, J.C., Chu, W.H., Xiang, X., Chen, F., Wang, C.S., Feng, H., Lin, J.K., 2014. Curcumin attenuates acute inflammatory injury by inhibiting the TLR4/MyD88/NF-kappaB signaling pathway in experimental traumatic brain injury. J Neuroinflammation 11, 59.

